# Transpeptidase PBP2 governs initial localization and activity of major cell-wall synthesis machinery in *Escherichia coli*

**DOI:** 10.1101/716407

**Authors:** Eva Wollrab, Gizem Özbaykal, Antoine Vigouroux, Baptiste Cordier, Francois Simon, Thibault Chaze, Mariette Matondo, Sven van Teeffelen

## Abstract

Bacterial shape is physically determined by the peptidoglycan cell wall. The cell-wall-synthesis machinery responsible for rod shape in *Escherichia coli* is the processive ‘Rod complex’. Previously, cytoplasmic MreB filaments were thought to govern formation and localization of Rod complexes based on local cell-envelope curvature. However, using single-particle tracking of the transpeptidase PBP2, we found strong evidence that PBP2 initiates new Rod complexes by binding to a substrate different from MreB or any known Rod-complex component. This substrate is likely the cell wall. Consistently, we found only weak correlations between MreB and envelope curvature in the cylindrical part of cells. Residual correlations do not require any curvature-based Rod-complex initiation but can be attributed to persistent rotational motion. Therefore, local cell-wall architecture likely provides the cue for PBP2 binding and subsequent Rod-complex initiation. We also found that PBP2 has a limiting role for Rod-complex activity, thus supporting its central role.

## Introduction

The peptidoglycan (PG) cell wall is the major load-bearing structure of the bacterial cell envelope and physically responsible for cell shape (Vollmer et al., 2008). Rod-like cell shape in *Escherichia coli* and other rod-like model organisms requires peptidoglycan synthesis by stable multi-enzyme ‘Rod complexes’ containing the transglycosylase RodA, the transpeptidase PBP2, the transmembrane protein RodZ, and the actin homolog MreB (Cho et al., 2016; Emami et al., 2017; Meeske et al., 2016; Morgenstein et al., 2015; Typas et al., 2012). All of these proteins move persistently around the cell circumference at similar speeds (Cho et al., 2016; Morgenstein et al., 2015; van Teeffelen et al., 2011), suggesting that these proteins stably associate for processive cell-wall insertion. Colocalization of MreB and RodZ (Alyahya et al., 2009; Bendezú et al., 2009; Morgenstein et al., 2015) supports this idea.

Other proteins (MreC, MreD, PBP1a, and PBP1b) are possibly also part of these complexes (Banzhaf et al., 2012; Cho et al., 2016; Contreras-Martel et al., 2017; Kruse et al., 2004; Morgenstein et al., 2015). MreC activates PBP2 (Contreras-Martel et al., 2017; Rohs et al., 2018). However, the shape defect of a *mreCD* deletion is partially suppressed by a hyperactive PBP2 point mutant (Rohs et al., 2018), suggesting that neither MreC nor MreD are strictly necessary for Rod-complex assembly or function. The bi-functional class-A penicillin-binding proteins PBP1a and PBP1b interact with PBP2 and RodZ, respectively (Banzhaf et al., 2012; Morgenstein et al., 2015), and PBP2 activates PBP1a glycosyltransferase activity *in vitro* (Banzhaf et al., 2012). However, Rod-complex rotational motion is independent of class-A PBP activity (Cho et al., 2016). Furthermore, single-molecule tracking suggests that any possible association of PBP1a or PBP1b with the Rod complex is short lived (Cho et al., 2016). Similar to *mreCD*, a *rodZ* deletion can also be suppressed by point mutations in PBP2, RodA, or MreB (Shiomi et al., 2008). Summarizing, it emerges, that RodA, PBP2, and MreB form the core of the Rod complex (Rohs et al., 2018). On the contrary, the determinants of Rod-complex spatial distribution and activity, which are ultimately responsible for cell shape, remain less well understood.

MreB filaments are intrinsically curved (Hussain et al., 2018; Salje et al., 2011). This curvature likely stabilizes the uniform circumferential or near-circumferential orientation of MreB filaments in the cylindrical parts of rod-shaped cells (Billaudeau et al., 2019; Hussain et al., 2018; Olshausen et al., 2013; Ouzounov et al., 2016; Wang and Wingreen, 2013), while uniform orientation is lost in aberrantly shaped cells (Hussain et al., 2018). MreB filaments are therefore likely responsible for the stable near-circumferential orientation of both Rod complex movement (Errington, 2015; Hussain et al., 2018) and the cell wall (Hussain et al., 2018; Wang et al., 2012). Computational simulations suggest that circumferential organization of the cell wall, in turn, contributes to mechanical integrity (Morgenstein et al., 2015; van Teeffelen et al., 2011) and maintenance of rod-like cell shape (Nguyen et al., 2015).

Previously, it has been suggested that MreB filaments provide a platform that recruits other Rod-complex components to the site of future cell-wall synthesis (Errington, 2015; Shi et al., 2018; Surovtsev and Jacobs-Wagner, 2018). Accordingly, MreB filaments might be responsible for the initial localization of Rod complexes. Ursell et al. and others suggested that MreB filaments are attracted to sites of specific two-dimensional cell-envelope curvature (Billings et al., 2014; Shi et al., 2018; Ursell et al., 2014) based on mechanical properties of MreB filaments and RodZ-MreB interactions (Bratton et al., 2018; Colavin et al., 2018). However, correlations could also come about indirectly, for example through a curvature-independent depletion of MreB from highly curved cell poles (Kawazura et al., 2017) or through persistent motion (Hussain et al., 2018; Wong et al., 2017, 2019). Therefore, the initial localization of Rod complexes could in principle be governed by factors different from MreB. We thus wondered, whether the cell wall itself could provide a local cue for the initiation of Rod complexes, independently of cell-envelope curvature. Such a local cue would have to be sensed by a protein with a periplasmic domain that can possibly bind the cell wall.

An obvious candidate is the transpeptidase PBP2. For its cross-linking activity PBP2 must bring together donor peptides on nascent glycan strands and acceptor peptides in the cell wall. Binding of PBP2 to the existing cell wall could therefore provide an alternative mechanism of Rod-complex initiation. In support of this hypothesis, a PBP2(L61R) mutant shows increased cell-wall synthetic activity and affects the distribution of MreB-actin filament length (Rohs et al., 2018). Further support comes from localization studies in *Caulobacter crescentus*: There, the spatial distributions of PBP2 and MreB only partially overlap, which is compatible with the hypothesis that PBP2 finds sites for activity independently of MreB (Dye et al., 2005; Hocking et al., 2012).

Here, we used single-molecule tracking to demonstrate that diffusive PBP2 molecules stably bind to an immobile and non-saturating component in the cell envelope. Bound molecules transition between immobile and persistently moving states, the latter depending on Rod-complex activity. Interestingly, MreB filaments are not the binding substrate for PBP2, nor does MreB determine locations of PBP2 binding. Other known Rod-complex components are likely also not required. A high degree of residual persistent motion upon RodA repression suggests that PBP1a or a different transglycosylase can contribute to processive Rod-complex activity (Banzhaf et al., 2012).

Our major observation let us conclude that PBP2 determines the initial localization of newly forming Rod complexes, possibly by binding to the cell wall directly. In support of this conclusion, we found that MreB filaments are likely not recruited to regions of particular cell-envelope curvature in filamentous and normal cells, contrary to (Ursell et al., 2014). Specifically, we found only weak MreB-curvature correlations once cell poles were excluded from the analysis. Residual correlations were attributed to weak spontaneous cell bending, suggesting that they are caused by persistent rotational motion (Wong et al., 2017). Finally, we also found that fast diffusing molecules cannot contribute to processive Rod-complex activity due to limitations of diffusion, contrary to (Lee et al., 2014).

## Results

### PBP2 enzymes can be quantitatively separated into diffusive and bound fractions

To study the role of PBP2 for the formation of Rod complexes we characterized its different states of motion, which are potentially representative of different states of substrate binding and activity. We imaged a functional, N-terminal protein fusion of the photo-activatable fluorescent protein PAmCherry to PBP2 (Lee et al., 2014). The fusion is expressed from the native *mrdA* locus at a level similar to the wild-type protein according to quantitative mass spectrometry (Fig. 1–SI Fig. 1D) and Bocillin labeling experiments [Fig. 1–SI Fig. 1A and also (Lee et al., 2014)]. The strain carrying the fusion maintains rod-like cell shape with only slight deviations of average cell diameter and length (Fig. 1–SI Fig. 1B) and does not show any growth defect (Fig. 1–SI Fig. 1C) (Lee et al., 2014).

**Figure 1.**
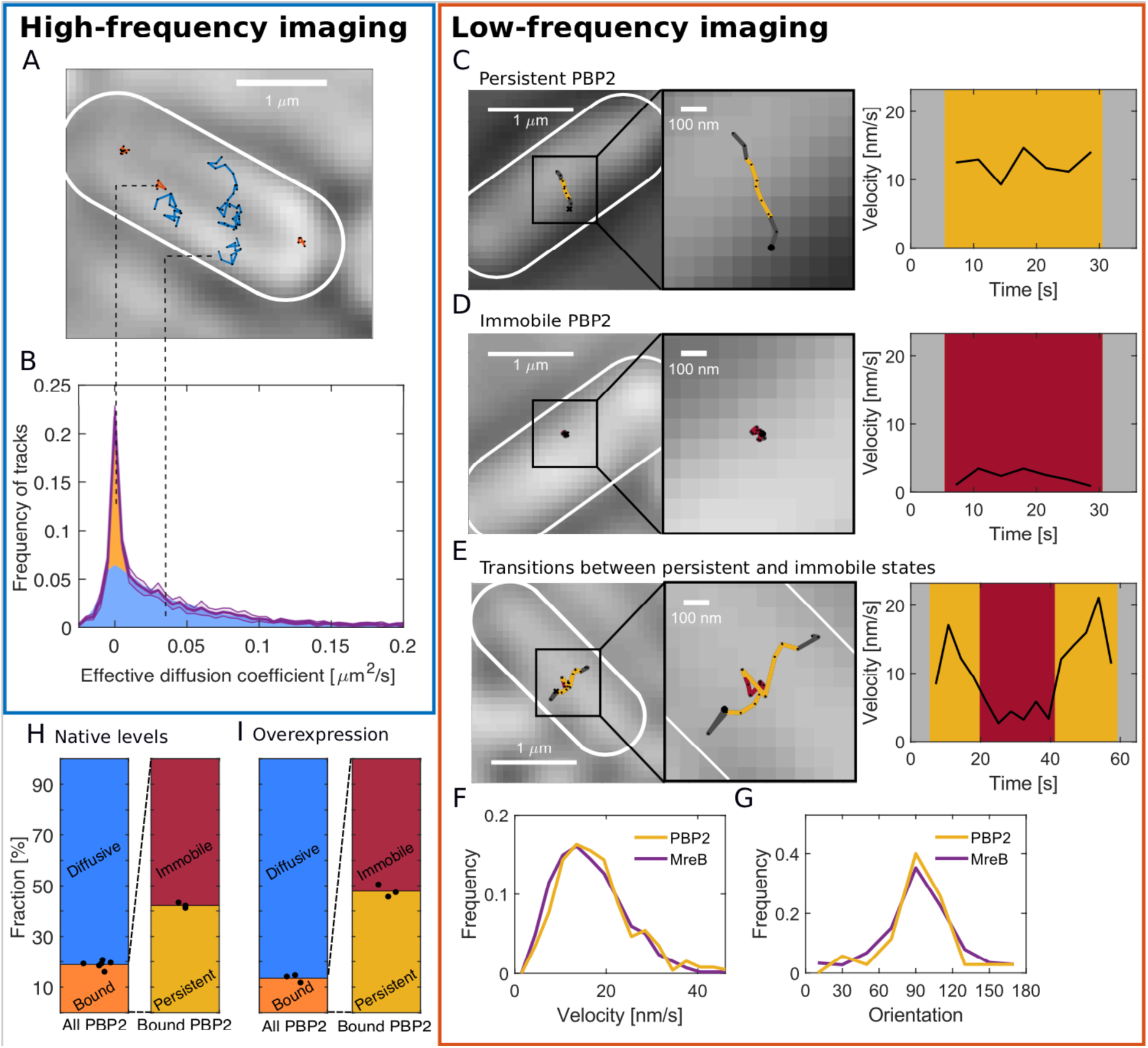
PBP2 molecules reside in diffusive, immobile, or persistently moving states. **(A)** Representative trajectories of PAmCherry-PBP2 molecules (TKL130) obtained by high-frequency imaging (time interval 60 ms) reveals diffusive (blue) and bound (orange) molecules. **(B)** Probability distribution of single-molecule effective diffusion coefficients (purple) and fit to a two-state diffusion model. 81% of PBP2 move diffusively with *D* = 0.039 μm^2^/s (blue region) while 19% are immobile (orange region). The shaded area in magenta indicates standard deviation between replicates. **(C-E)** Low-frequency imaging (3.6 s with 1 s exposure time) reveals that bound PBP2 molecules are either persistently moving **(C)** or immobile **(D)**, according to the instantaneous PBP2 velocity. PBP2 molecules show transitions between persistent and immobile states **(E)**. **(F-G)** Persistently moving PBP2 and MreB filaments show similar speeds **(F)** and orientations of motion (orientation measured with respect to the cell centerline) **(G)**. **(H-I)** Average fractions of bound, diffusive, persistently moving, and immobile PAmCherry-PBP2 at native levels (TKL130) **(H)** or if overexpressed (TKL130/pKC128) **(I)**. Dots show biological replicates.

We obtained single-molecule tracks by single-particle tracking *P*hoto*A*ctivatable *L*ocalization *M*icroscopy (sptPALM) (Manley et al., 2008) in total internal reflection fluorescence (TIRF) mode, which restricts the observation to the bottom part of the cell. We first imaged PBP2 molecules at high frequency (intervals of 60 ms). We found both spatially extended trajectories, corresponding to fast diffusing molecules, and trajectories that appeared as localized, corresponding to immobile or slowly moving molecules (Fig. 1A, Fig. 1–Movie 1). We confirmed the presence of two distinct fractions of diffusing and localized molecules based on the distribution of single-track effective diffusion constants (Fig. 1B). More specifically, the experimental distribution was fit to the prediction from a two-state diffusion model, which contains as a special case a diffusing and an immobile population (Fig. 1B; Fig. 1–SI Fig. 2). We found a population with diffusion constant *D*_1_ = 0.039 ± 0.002 μm^2^/s containing 19 ± 2% of all enzymes, and a second population with zero diffusion constant containing 81 ± 2% of all molecules.

The relative sizes of the two populations are only weakly affected by PBP2 expression level: Upon overexpressed by about three- to six-fold (Fig. 1–SI 1D) using the vector pKC128 that expresses PAmCherry-PBP2 from the native *mrdAB* promoter (Lee et al., 2014) the fraction of molecules with near-zero diffusion constant decreased only mildly from 20% to 15% (Fig. 1H-I and Fig. 1–SI Fig. 3).

While the diffusing molecules are likely enzymatically inactive and possibly searching for new sites of cell-wall insertion (see further down), the localized molecules are therefore likely bound to an immobile and non-saturated substrate or part of slowly moving Rod complexes with anticipated speeds of 10-40 nm/s (Cho et al., 2016; van Teeffelen et al., 2011).

### Bound PBP2 molecules are either persistently moving or immobile

To test whether all or part of the bound molecules were moving persistently we imaged PBP2 molecules at low frequency, taking images with an exposure time of 1 s and intervals of 3.6 s. The long exposure time effectively smears out the fluorescence of fast diffusing molecules, allowing us to detect the positions of individual bound molecules. Using this protocol, we found molecules that moved persistently, were immobile, or showed transitions between these two states (Fig. 1C-E, Fig. 1–Movie 2, and Fig. 1–SI Fig. 4).

Persistently moving molecules showed similar distributions of speed and orientation as a functional msfGFP-MreB fusion (Ouzounov et al., 2016) expressed from the native *mreB* locus (Fig. 1F, G) (Cho et al., 2016)], in agreement with previous measurements (Cho et al., 2016; van Teeffelen et al., 2011). For accurate velocity measurements, we obtained these results from movies acquired with a shorter time interval of 1 s. Straight tracks representing persistently moving molecules were selected based on the mean squared displacements (MSD) (Fig. 1–SI Fig. 5).

Because PBP2 molecules show transitions between different states in single trajectories, we quantified immobile and persistent states locally in time. Specifically, we classified motion states using a threshold on the mean velocity during four consecutive time steps in movies acquired with 3.6 s interval (Fig. 1C-E). Window size and velocity threshold (8 nm/s) were chosen based on computationally simulated tracks (Fig. 1–SI Fig. 6A-C). In confirmation of our two-state model, we found good agreement between the average MSD obtained from experiment and simulation for immobile and persistent segments, respectively (Fig. 1–SI Fig. 6D-F). Other states of motion are therefore likely not present. Using this criterion we found a persistent fraction of 42.2 ± 1.1 % of all bound molecules, while 57.8 ± 1.1 % remained immobile (Fig. 1H).

Upon overexpression of PAmCherry-PBP2 as above, we found that the persistent fraction remained nearly constant (Fig. 1H). This finding suggests that PBP2 limits the number of active Rod complexes. This viewpoint is consistent with the recent report that a hyperactive PBP2 point mutant (L61R) increased the overall amount of active Rod complexes (Rohs et al., 2018). We will come back to this mutant below.

### msfGFP-PBP2 fusion confirms findings and demonstrates increased PBP2 binding upon PBP2 depletion

We confirmed our findings using a strain that carries a functional msfGFP-PBP2 fusion under IPTG-inducible control as the sole copy of PBP2 (Cho et al., 2016). We chose the induction level (25 uM IPTG) based on measurements of cell diameter (Fig. 1–SI Fig. 7A). Noteworthy, quantitative mass spectrometry showed levels of msfGFP-PBP2 to be about three-fold higher than native PBP2 in the wild type (Fig. 1–SI Fig. 7B), while near-wild-type levels observed at lower induction (5 uM IPTG) led to loss of rod shape at long times (Fig. 1–SI Fig. 7A). This discrepancy might be a consequence of reduced enzymatic efficiency of the msfGFP fusion or simply due to the more noisy expression from the inducible promoter.

Tracking the msfGFP-PBP2 at high frequency after initial pre-bleaching, we found a similar fraction of bound PBP2 molecules (23.2 ± 9.2 %) (Fig. 1–SI Fig. 7C, Fig. 1–Movie 3). Similarly to PAmCherry-PBP2, the bound fraction was reduced only slightly upon two-fold increase of protein levels (18 ± 6.1 %), supporting our previous conclusion that the target of PBP2 binding is non-saturated.

Among the bound molecules we found a fraction of 80 ± 10 % persistently moving molecules (Fig. 1–SI Fig. 8A, Fig. 1–Movie 4), which is significantly higher than in the case of the PAmCherry fusion. However, similarly to the PAmCherry fusion, the persistent fraction remained nearly constant over the 6-fold range of expression levels, only showing a mild drop at the highest induction level of 100 uM IPTG (Fig. 1–SI Fig. 8A). These results therefore confirm that the binding target of PBP2 is non-limiting and that PBP2 actively limits Rod-complex activity.

Interestingly, the bound fraction increased almost two-fold after long times of low expression levels (5 uM) (Fig. 1–SI Fig. 9B), already before rod shape had been lost (Fig. 1–SI Fig. 9A). This observation suggests a potential feedback mechanism between cell-wall architecture due to reduced Rod-complex activity and PBP2 binding. We will come back to this point further down.

### A hyperactive PBP2 mutant (L61R) shows increased binding at low expression

Initially intended as a further test of our method, we also tracked an msfGFP fusion to the PBP2 point mutant (L61R) recently characterized by (Rohs et al., 2018). Based on slow-frequency imaging, they reported that the number of bound msfGFP-PBP2(L61R) was about two-fold increased as compared to msfGFP-PBP2 (Rohs et al., 2018). We confirmed this finding quantitatively (Fig. 1–SI Fig. 7C). However, we also found that the bound fraction was high only at low protein expression (5 uM IPTG), while it was equal to the bound fraction of the wild-type protein at higher expression (25 uM IPTG). Furthermore, the persistent fraction was reduced in comparison to the wild-type protein (Fig. 1–SI Fig. 8A). Therefore, the mutant shows higher activity at low expression as previously reported but reduced activity at higher expression.

### PBP2 molecules show long persistent runs

The classification into different motion states at the sub-trajectory level allowed us to extract average transition rates between immobile and persistently moving states, *k*_ip_ and *k*_pi_, respectively (Fig. 2A-B). Depending on the fluorescent-protein fusion, we found values of *k*_ip_ between 0.015-0.06 s^-1^, and of *k*_pi_ between 0.006-0.02 s^-1^. The msfGFP-PBP2 fusion shows less frequent arrests and faster transitions from immobile to persistently moving states, in agreement with its higher fraction of moving molecules (Fig. 1–SI Fig. 8A).

**Figure 2.**
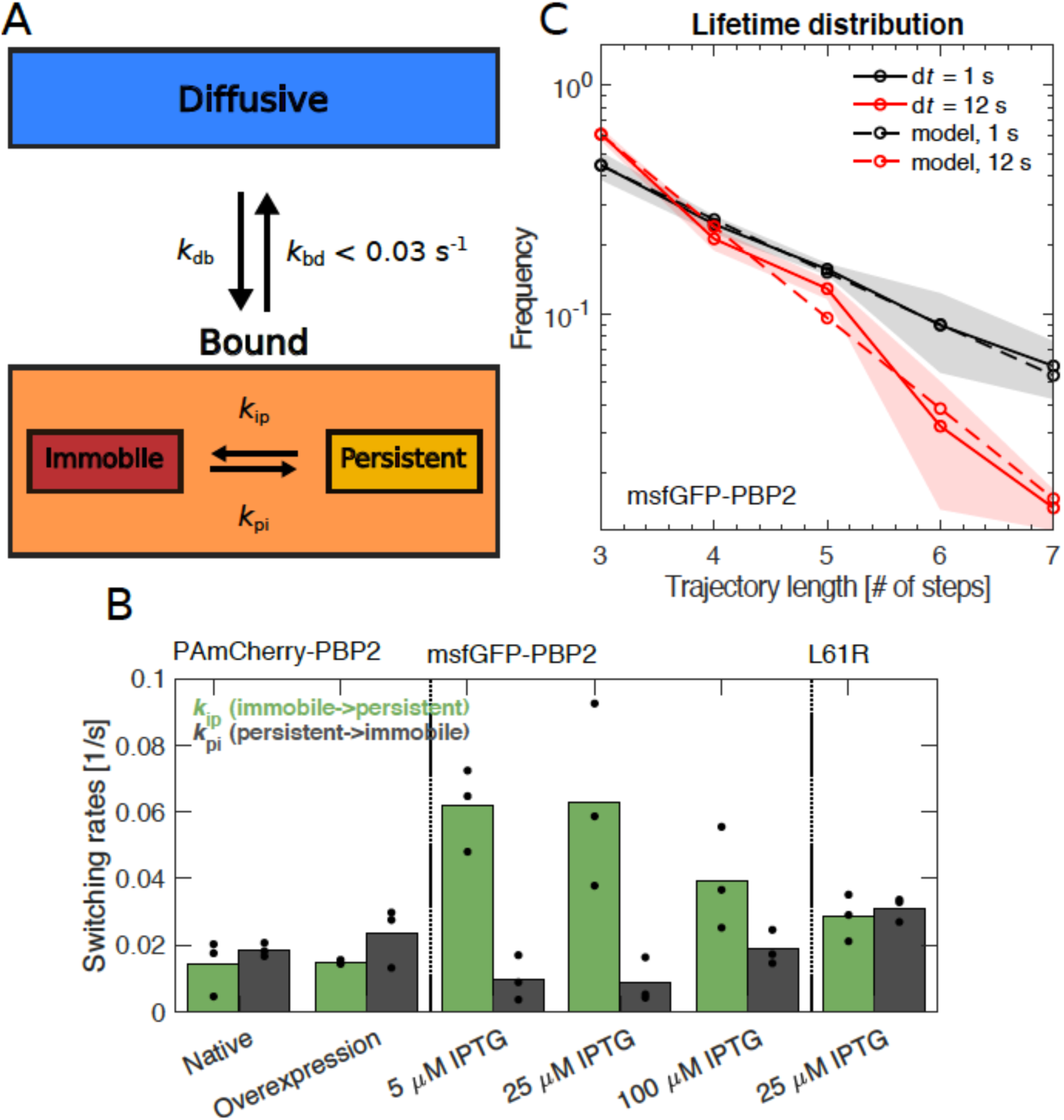
PBP2 molecules transition between different dynamic states. **(A)** Diagram illustrating transition rates measured between different motion states. **(B)** Transition rates between immobile and persistently moving states for different protein fusions and expression levels. Circles: biological replicates. **(C)** Fluorescence-lifetime distributions of msfGFP-PBP2 trajectories with imaging intervals of 1 s (black solid line) and 12 s (red solid line). Dashed lines represent a joint fit of the two curves to a model of photobleaching and bleaching-independent track termination, the latter comprising unbinding and persistently molecules leaving the TIR field of view (bleaching probability per frame *p*_b_ = 0.39 ± 0.08, apparent track termination rate *k*_a_ = 0.035 ± 0.007 s^-1^). Based on a model for persistent motion, we obtained an upper limit of the unbinding rate of *k*_bd_ < 0.03/s (Fig. 2–SI Fig. 1). Shaded region: Standard deviation between replicates.

A persistent run of PBP2 terminates either due to an arrest of PBP2 (persistent-to-immobile transition) or due to an unbinding event (persistent-to-diffusive transition). As an upper bound of the unbinding rate, we measured the transition rate from the aggregate bound state (persistent and immobile states) to the diffusive state, *k*_bd_ (Fig. 1A). Specifically, we acquired distributions of track lengths for two different imaging intervals of 1 s and 12 s (Fig. 2C), using the msfGFP-PBP2 fusion for the higher number of tracks obtained. Track length is limited by bleaching, unbinding, and persistent molecules leaving the field of view. The latter two processes are responsible for the shorter track lengths observed for d*t* = 12 s. Taking all three processes into account in computational simulations, we obtained an upper limit of the unbinding rate of *k*_bd_ < 0.03 s^-1^, corresponding to a minimum average lifetime of 30 s (Fig. 2– SI Fig. 1).

The rates *k*_pi_ and *k*_bd_ then yield the possible ranges of average run lengths of persistently moving molecules between 0.3-1.6 um, depending on protein fusion and exact unbinding rate. This range is compatible with long tracks of MreB motion observed previously (van Teeffelen et al., 2011).

### PBP2 spatial pattern and bound fraction are independent of MreB cytoskeleton

MreB is often regarded as a hub for Rod-complex components (Errington, 2015; Shi et al., 2018; Surovtsev and Jacobs-Wagner, 2018). To determine whether MreB is the substrate of PBP2 binding we treated cells with the putative MreB-polymerization inhibitor A22 (Gitai, 2005). A22 treatment strongly reduced both number and size of MreB-msfGFP spots observed on the two-dimensional cell contour using epi-fluorescence microscopy (Fig. 3A, Fig. 3–SI Fig. 1A) and abolished rotational motion (Fig. 3–Movies 1-2).

**Figure 3.**
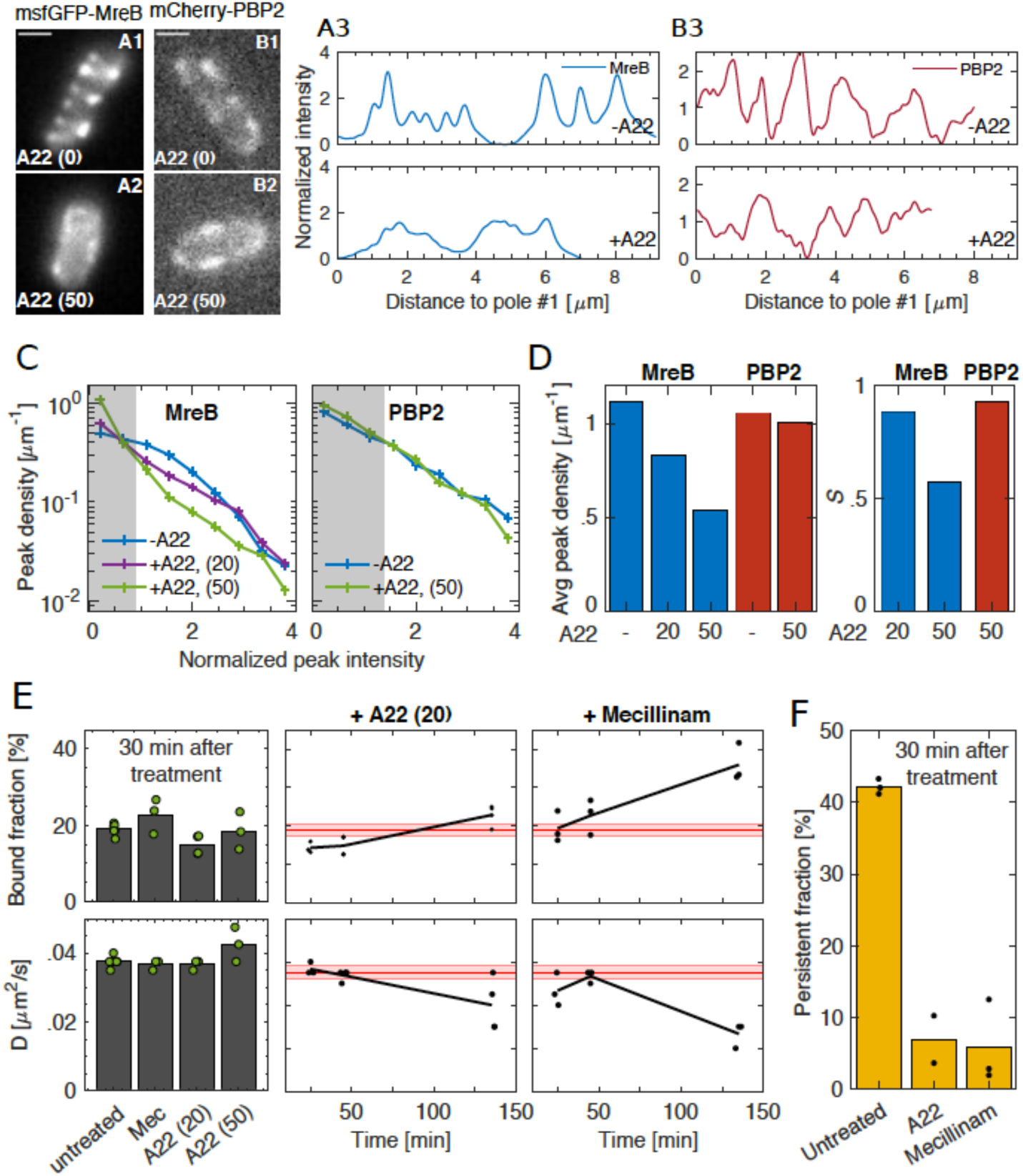
Spatial distribution and magnitude of PBP2 bound fraction are independent of MreB cytoskeleton. **(A-B)** A22 (30 min; 50 μg/ml) visibly reduces peak number and intensity of MreB-msfGFP **(A)** but not of mCherry-PBP2 **(B)** on cell boundaries (image exposure time 1 s). Scale bar 1 μm. **(C)** Peak density on the cell boundary [1/μm] as function of peak intensity for two A22 concentrations (20, 50 μg/ml). Intensities are normalized by median peak intensity in untreated cells. Gray regions: peaks within noise floor. **(D) Left**. Density of all peaks above noise floor in (A). **Right**. Fold-change *S* of the amount of molecules found inside peaks between untreated and A22-treated conditions. **(E) Left**. Bound fraction and diffusion constant of PAmCherry-PBP2 30 minutes after drug treatment with mecillinam (labeled ‘Mec’, 100 μg/ml) or A22 (20 or 50 μg/ml). **Right**. Bound fraction and diffusion constant over time after treatment with A22 (20 μg/ml) or mecillinam (100 μg/ml). Dots indicate biological replicates. Red lines and shaded areas: Average values and standard deviations between replicates from untreated cells. **(F)** Persistent fractions corresponding to the 30 min time point in (E).

To test whether MreB filaments were required for PBP2 binding we first imaged the spatial distribution of an mCherry-PBP2 fusion on the cell contour in untreated or A22-treated conditions, in the same manner as we imaged MreB-msfGFP (Fig. 3B, Fig. 3–SI Fig. 1B). We found that mCherry-PBP2 showed a spotty pattern. Since these images were taken with a long exposure time (1s), each spot likely represents multiple bound molecules. Different from MreB-msfGFP, the spotty pattern of mCherry-PBP2 was not affected by A22 treatment (Fig. 3C) – a strong indication that MreB filaments are not the binding substrate for PBP2.

Next, we measured bound and persistent fractions of PBP2 molecules before and after A22 treatment using single-molecule tracking in TIRF. The bound fraction remained close to the value of untreated cells (Fig. 3E). Yet, the apparent fraction of persistently moving molecules among all bound PBP2 enzymes nearly vanished (Fig. 3F). This is consistent with the arrest of MreB rotation (Fig. 3–Movie 2) and with previous bulk measurements of Rod-complex activity (Uehara and Park, 2008). To follow the bound fraction during two mass-doubling times (Fig. 3E), we used an intermediate concentration of A22 (20 uM), which did not affect growth (Fig. 3–SI Fig. 2A). Together, our findings suggest that MreB polymers are neither the substrate of PBP2 binding nor do they affect the rate of PBP2 binding and unbinding.

MreB depolymerization did also not elicit a rapid change of the diffusion constant of the freely diffusing molecules (Fig. 3E), suggesting that MreB does not significantly constrain the movement of diffusive PBP2 molecules to an area close to the MreB cytoskeleton (Strahl et al., 2014). Therefore, if PBP2 binds its substrate from the diffusive state, the locations of PBP2 binding are also independent of MreB.

A22 treatment already demonstrates that PBP2 binding is independent of Rod-complex activity. We confirmed this finding using the PBP2-targeting beta-lactam Mecillinam, which binds covalently to the active site (Spratt, 1975). Mecillinam did not cause a reduction of the bound fraction (Fig. 3E), demonstrating that PBP2 binds its binding target with a moiety different from its active site.

At long times of treatment with Mecillinam or A22 (120 min) the bound fraction increased and the diffusion constant decreased (Fig. 3E), which coincides and is potentially caused by the loss of normal cell-wall architecture during loss of rod-like cell shape (Fig. 3–SI Fig. 2C), similar to the increase of the bound fraction at sustained low induction levels of msfGFP-PBP2 (Fig. 1–SI Fig. 9B).

### PBP2 binds to its substrate at locations that are independent of MreB localization

To demonstrate that PBP2 binding was indeed independent of MreB filaments or Rod-complex activity as suggested by Fig. 3 we still needed to show that PBP2 molecules interchange between diffusive and bound states during A22 treatment. We already found a low upper bound for the transition rate from bound to diffusive states in non-treated cells (*k*_bd_ < 0.03 s^-1^) (Fig. 2C), and we expect inverse transitions to occur even more rarely. We therefore used a variant of Fluorescence Recovery After Photobleaching (FRAP) (Fig. 4A) termed Bound-Molecule FRAP: Instead of measuring fluorescence intensity we measure the bound fraction at different time points after bleaching almost all molecules at the bottom of the TIR field of view. Right after bleaching, the bound fraction dropped significantly (Fig. 4B), suggesting that fast diffusing molecules re-entered the observation window within less than half a minute but did not quickly bind their substrate. Within about 2-4 min the bound fraction recovered, yielding a transition rate from diffusive to bound states of *k*_db_ = (4.5 ± 2)*10^−3^ s^-1^. The same experiment in non-treated cells did not reveal recovery of the bound fraction (Fig. 4–SI Fig. 1), presumably because bound molecules left the field of view through persistent motion.

**Figure 4.**
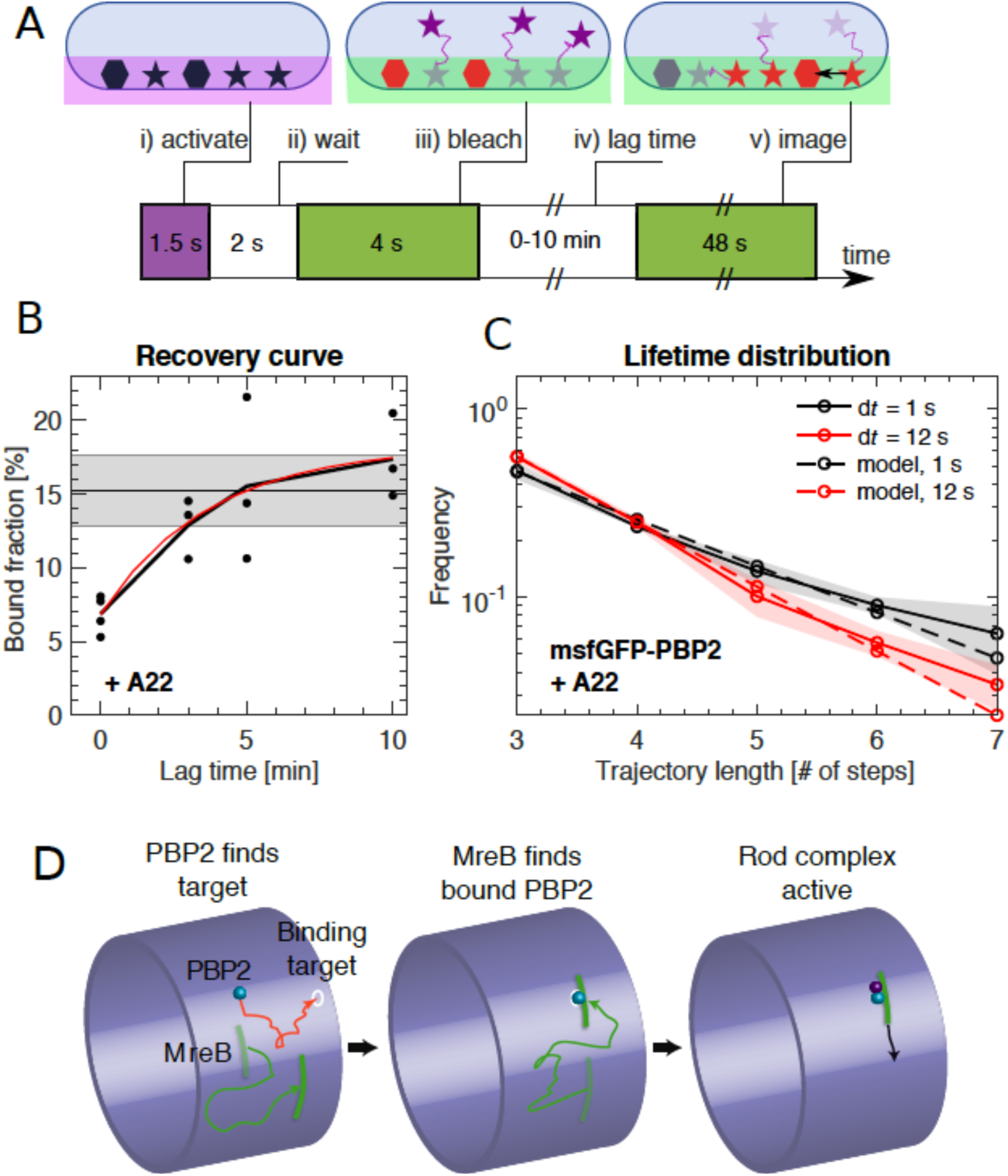
PBP2 slowly transitions between diffusive and bound states. **(A)** Bound-Molecule-FRAP reveals rate of PAmCherry-PBP2 binding *k*_db_: i) Diffusive (stars) and bound (hexagons) molecules are activated at bottom of cell through TIR illumination. ii) Most activated diffusive molecules (purple) leave the field of view. iii) Remaining molecules are bleached (red). iv) Activated diffusive molecules partially return into the field of view, where they can bind (black arrow). v) Measurement of bound fraction. **(B)** Bound fraction according to (A) at different lag times. Black horizontal line and shaded area: bound fraction without bleaching and standard deviation from replicates. An exponential fit in the form *b*(*t*) = *a*_1_ - *a*_2_ exp[-*k*_db_ *t*] (red line) yields binding rate *k*_db_ = (4.5 ± 2)*10^−3^ s^-1^. **(C)** Fluorescence-lifetime distributions of msfGFP-PBP2 tracks in A22-treated cells with imaging intervals of 1 s (black solid line) and 12 s (red solid line) yields unbinding rate *k*_bd_ = 0.02 ± 0.01 s^-1^. **(D)** Cartoon of suggested Rod-complex initiation through PBP2: PBP2 (blue) binds to a target site in the cell envelope (white circle) independently of MreB filaments or PBP2 activity. PBP2 then recruits an MreB filament through diffusion and capture (green) or through nucleation, and also recruits other rod-complex components (magenta).

Since the bound and diffusive fractions did not change by more than 10% after A22 treatment, transition rates from the diffusive into the bound states must be matched by reverse transitions from the bound to the diffusive state with a rate *k*_bd_ = *k*_db_(1-*b*)/*b* = 0.02 ± 0.01 s^-1^, where *b* is the bound fraction. We confirmed this expectation through independent lifetime measurements of A22-treated cells, similar to those in Fig. 2 (Fig. 4C), yielding *k*_bd_ = 0.018 ± 0.01 s^-1^. Since bound fractions are almost identical for untreated and A22-treated cells, we reasoned that binding and unbinding rates *k*_bd_ and *k*_db_ are likely also the same in both conditions.

The average lifetime of a bound molecule of about 1 min is 70-fold smaller than the cell doubling time of 70 min (Fig. 1–SI Fig. 1C). Therefore, almost all bound PBP2 molecules observed at any time have undergone multiple transitions from the diffusive to their current bound state. Together with the previous observation that free diffusion is not constrained by MreB filaments we can thus conclude that PBP2 binds its substrate at locations that are determined independently of MreB filaments.

### PBP2 determines the location of new Rod complexes

Two qualitatively different scenarios for the formation of an active Rod complex are conceivable, once PBP2 has bound to a location in the cell envelope: First, MreB filaments could join PBP2 *after* PBP2 binding (through diffusion and capture or through nucleation; Fig. 4D). Alternatively, by chance, MreB filaments could already be present at the PBP2-binding site *prior* to PBP2 binding (however, without actively influencing the sites of PBP2 binding itself, as shown in the previous paragraph).

Since MreB filaments cover only a small area fraction of the cytoplasmic membrane [about 2% according to a conservative estimate (Methods)], the latter scenario would require that a single PBP2 molecule bound and unbound unsuccessfully more than 50 times before finding an MreB filament by chance. With the binding and unbinding rates measured it would take an initially diffusing molecule about 200 minutes to become part of an active rod complex, which makes this scenario highly unlikely. Therefore, we suggest that PBP2 binds to a site in the cell envelope and then recruits MreB filaments (Fig. 4D). Thus, PBP2 determines the locations of newly forming Rod complexes.

### PBP2-binding substrate is none of the known Rod-complex components but likely the cell wall

Proteins different from MreB could still provide the binding substrate or be required for binding – specifically the putative Rod-complex components MreC, MreD, RodA, RodZ, and PBP1a. We therefore constructed depletion strains for RodA, RodZ, or MreCD in a background strain expressing either native levels of PAmCherry-PBP2 (for RodZ, MreCD) or overexpressing PAmCherry-PBP2 (for RodA). Without repression, all of the strains showed normal growth rate (Fig. 5–SI Fig. 1), cell shape (Fig. 5A, Fig. 5–SI Fig. 2), bound fractions (Fig. 5C), and persistent fractions (Fig. 5D). Only the RodZ-depletion strain showed slightly higher bound and persistent fractions.

**Figure 5.**
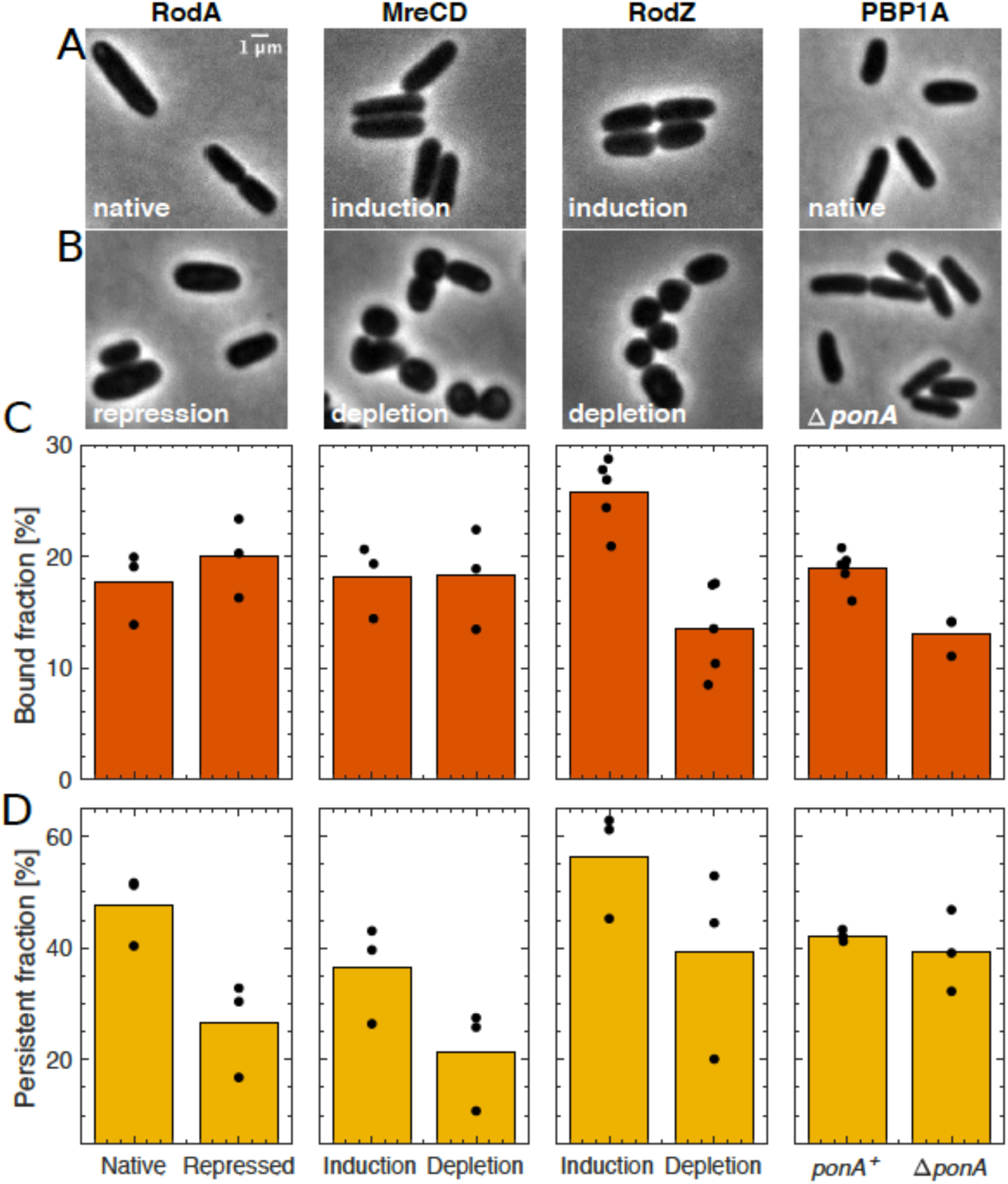
Depletion of known rod-complex components only weakly affects PBP2 binding. **(A-B)** Cell shape upon near-native expression **(A)** or long-time depletion **(B)** of RodA, MreCD, RodZ, or PBP1a. RodA was repressed for 9h through CRISPRi against *mrdAB* operon (coding for PBP2 and RodA) in AV48/pKC128 [P_*mrdA*_::*PAmCherry-PBP2*]. Here, PBP2 was 2-5-fold overexpressed from plasmid pKC128 to avoid PBP2 depletion upon *mrdAB* repression. MreCD was depleted for 6h in TKL130 Δ*mreCD*/pFB121 [P_lac_::*mreCD*]. RodZ was depleted for 6h in TKL130 Δ*rodZ*/pFB290 [P_lac_::*rodZ*]. PBP1a is not essential and was deleted. In all cases except for PBP1A, cells loose rod-like cell shape. **(C-D)** Bound fractions **(C)** and persistent fractions **(D)** of PAmCherry-PBP2 upon expression or depletion of proteins indicated above (A).

Within 3-5 h after depletion, cells started to lose their rod-like shape (Fig. 5–SI Fig. 2). After growing cells for about two additional doubling times (according to OD), we measured bound and persistent fractions (Fig. 5C-D). At this point, repressed protein levels were reduced well below wildtype levels according to mass spectrometry (Fig. 5–SI Fig. 3A) and cell shape was severely perturbed (Fig. 5B, Fig. 5–SI Fig. 2) while growth rate remained unperturbed (Fig. 5–SI Fig.1). PBP2 levels remained close to native levels during all experiments except for RodA depletion, where PBP2 remained overexpressed (Fig. 5–SI Fig. 3B).

Upon depletion of either RodA or MreCD the bound fraction remained constant, while the persistent fraction markedly dropped. These findings suggest that RodA or MreCD are important for Rod-complex activity but that they are neither required for PBP2 binding nor do they affect the rates of PBP2 binding or unbinding.

Different from RodA and MreCD, depletion of RodZ led to a significant reduction of the bound fraction 6 h after starting depletion, while the persistent fraction remained high. Noteworthy, the reduction of persistent motion (the product of persistent and bound fractions) of about two-fold is compatible with independent measurements of the density of persistently moving MreB filaments (Dion et al., 2019). When measuring the bound fraction upon RodZ depletion over time, we found that the drop already occurred within about two hours (Fig. 5–SI Fig. 4), suggesting that RodZ might modulate the rates of PBP2 binding or unbinding. However, after 6 h of depletion, the bound fraction started to increase steadily, demonstrating that RodZ is not strictly required for PBP2 binding.

To test whether RodZ might influence the location of PBP2 binding we measured the degree of co-localization of mCherry-PBP2 and RodZ-GFP along the contour of cells expressing both fusions as sole copies of the respective proteins (Fig. 6A-C, Fig. 6–SI Fig. 1). PBP2 and RodZ did only show weak or no visible co-localization, which was also reflected by low Pearson’s correlation coefficients measured in single cells (Fig. 6D). In contrast, correlations between RodZ-GFP and MreB-mCherry were always high and positive (Fig. 6B-D, Fig. 6–SI Fig. 1) as expected (Alyahya et al., 2009; Morgenstein et al., 2015). As an additional test we also studied the density of RodZ-GFP peaks upon treatment with A22 (Fig. 6E-F). We found that the pattern of RodZ changed in a similar manner as the pattern of MreB-msfGFP (Fig. 3), while mCherry-PBP2 did not (Fig. 3), suggesting that the spatial pattern of RodZ depends on MreB (Bendezú et al., 2009), while the spatial pattern of PBP2 binding is independent of both MreB and RodZ.

**Figure 6.**
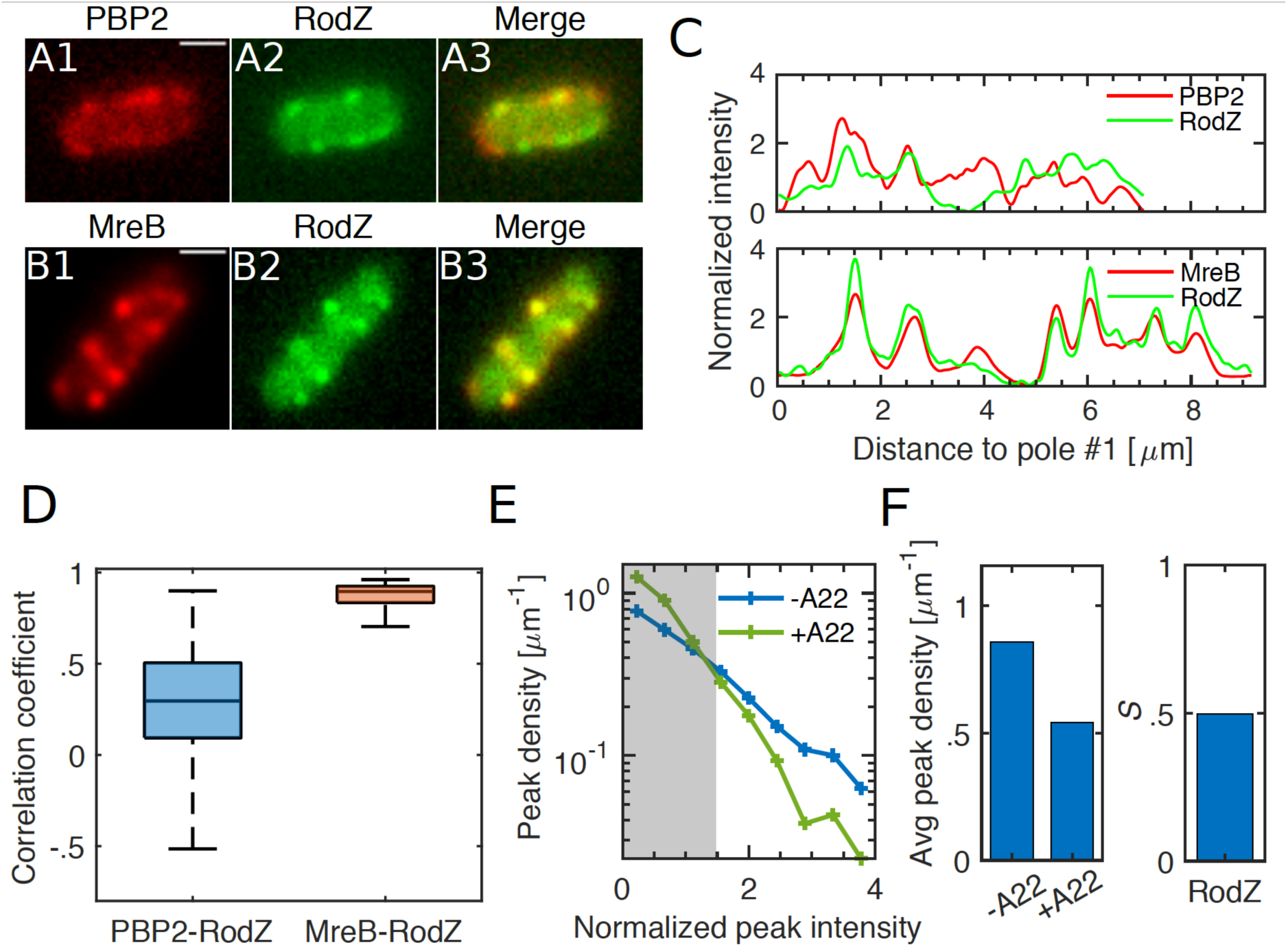
PBP2 localization does not depend on RodZ, and RodZ depends on MreB. **(A-B)** Fluorescence images of cells co-expressing mCherry-PBP2 and GFP-RodZ (Δ*rodZ mrdA*<>*mCherry-mrdA* (P_lac_::*gfp-rodZ*)) **(A)** or MreB-mCherry and GFP-RodZ (Δ*rodZ mreB*<>*mreB-mCherry* (P_lac_::*gfp-rodZ*)) **(B)** **(C)** Fluorescence signals along contours of cells displayed in (A) (mCherry-PBP2, GFP-RodZ) and (C) (MreB-mCherry, GFP-RodZ). **(D)** Pearson correlation coefficients between PBP2- and RodZ signals (left) and between MreB-and RodZ signals (right). **(E-F)** Localization pattern of GFP-RodZ (expressed as sole fusion and sole copy of RodZ in wt Δ*rodZ* (P_lac_::*gfp-rodZ*)) upon A22 treatment analyzed as in Fig. 3C-D. **(E)** Peak density on the cell boundary as a function of peak intensity. **(F) Left:** Density of all peaks above noise floor in (E). **Right:** The fold-change *S* of the amount of molecules found inside peaks between untreated and A22-treated conditions. GFP-RodZ behaves as MreB-msfGFP and differently from mCherry-PBP2 (Fig. 3C-D).

Qualitatively similarly to the effect of *rodZ* depletion, a *ponA* deletion strain (*ponA* codes for PBP1a) (Fig. 5) did not eliminate PBP2 binding but showed a drop of the bound fraction. Therefore, PBP1a is not required for PBP2 binding but might aid PBP2 binding or stabilize bound PBP2.

Our experiments suggest that none of the putative Rod-complex components are required for PBP2 binding or influence the spatial pattern of PBP2 binding. We therefore reasoned that PBP2 binds to the cell wall directly. Further support for this viewpoint comes from the diffusive motion of PBP2 molecules. PBP2 diffusion is much slower than diffusion of similar-size membrane proteins (Kumar et al., 2010) or of a truncated version of PBP2 (Lee et al., 2014), suggesting that PBP2 might weakly bind the cell wall even during diffusion. We found that depletion of RodA, RodZ, and MreCD caused an additional decrease of the diffusion constant (Fig. 5–SI Fig. 5), similarly to long-term treatment with A22 or Mecillinam (Fig. 3E). In all these experiments Rod-complex activity is inhibited or reduced, which changes cell-wall architecture (Wang et al., 2012) and reduces the degree of cross-linking (Uehara and Park, 2008). We therefore reasoned that diffusion is likely governed by the physical interactions between PBP2 and the cell wall (Lee et al., 2014), supporting the idea that PBP2 can bind the cell wall directly, through a domain that is different from its enzymatically active site.

### MreB-curvature correlations are likely the result of persistent motion

Previously, Ursell et al. observed that MreB filaments were excluded from regions of positive Gaussian cell-envelope curvature such as found at the cell poles, while MreB was enriched in regions of negative Gaussian curvature as found at the inner sides of bent cells (Ursell et al., 2014). They concluded that the locations of Rod-complex activity are determined by MreB filaments preferentially localizing to sites of negative Gaussian curvature in rod-shaped cells (Ursell et al., 2014). This conclusion is in contradiction to our finding that PBP2 is responsible for the initial localization of new Rod complexes in the cylindrical part of the cell. However, MreB-curvature correlations could also come about indirectly through persistent rotational motion (Hussain et al., 2018; Wong et al., 2017, 2019) or additional mechanisms of polar exclusion (Kawazura et al., 2017), without any curvature-based Rod-complex initiation. To resolve this conflict, we reinvestigated MreB-curvature correlations and their potentially different origin.

We followed a very similar approach to (Ursell et al., 2014). Specifically, we measured the spatial pattern of MreB-msfGFP (Ouzounov et al., 2016) on the two-dimensional cell contour (Fig. 7A,B) both in filamentous cells, through expression of the division inhibitor SulA (Bi and Lutkenhaus, 1993), and in non-filamentous cells growing on agarose pads under the microscope. We obtained the contour curvature of the cell from phase-contrast images using the Morphometrics cell segmentation tool (Ursell et al., 2017, 2014). In cylindrical regions of normally growing or filamentous cells with low variations of cell diameter *σ*, contour curvature *κ* is a good proxy for Gaussian curvature *G*, with *G* = 2 *k*/*σ*.

**Figure 7.**
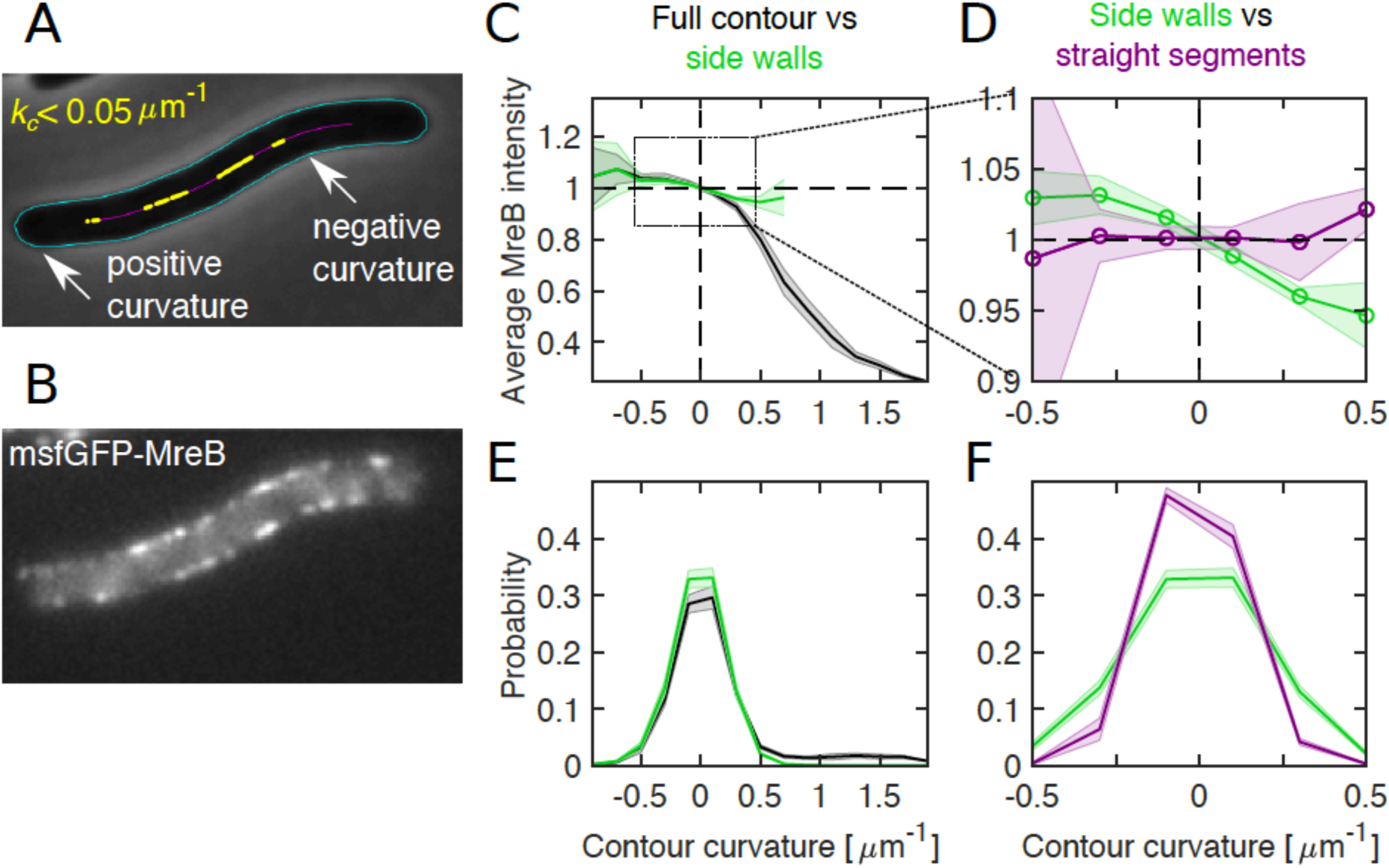
Differential MreB-curvature correlations in filamentous cells are due to cell poles and cell bending. **(A-B)** Phase-contrast image **(A)** and fluorescence intensity **(B)** of a representative filamentous *E. coli* expressing MreB-msfGFP and SulA (NO53/pDB192). Contours (cyan) are obtained by computational cell segmentation. Positive contour curvature is found at cell poles, bulges, and outer parts of spontaneously bent regions, while negative curvature is found at indentations and inner parts of bent regions. Straight cell segments (yellow) are defined as regions where the curvature of the spatially averaged centerline (magenta) is smaller than 0.05 μm^-1^. **(C-D)** Normalized average MreB intensity as a function of local contour curvature. Comparison between correlations obtained from full contours (black) and side walls (green) **(C)** and from side walls (green) and straight cell segments (magenta) **(D)**. **(E-F)** Distributions of contour-curvature values corresponding to correlation plots in (C-D). Shaded region: Standard deviation between replicates.

First, we measured the enrichment of MreB intensity at the cell contour as a function of local contour curvature, just like (Ursell et al., 2014) (Fig. 7C for filamentous cells; Fig. 7–SI Fig. 1 for wild-type cells). In quantitative agreement with their data we found enrichment of MreB at negative curvature and depletion at positive curvatures, as present at cell poles.

We then constrained our analysis to the cylindrical part of the cells and found that correlations were reduced by about five-fold at positive curvature values (Fig. 7C). These findings demonstrate that the pattern of MreB localization is not simply a function of contour or Gaussian curvature. Instead, curvature correlations are qualitatively different at different parts of the cell and dominated by cell poles. On the contrary, correlations between MreB and contour curvature in the cylindrical part of the cell are weak.

In previous work by some of us (Wong et al., 2017) we found small but significant correlations between MreB and cell-centerline curvature in mechanically bent cells. We therefore wondered whether residual correlations between MreB and contour curvature in the cylindrical part of the cell observed here were dominated by weak cell bending (Fig. 7A) likely caused by cells attaching to the glass surface (Duvernoy et al., 2018). To that end we restricted our analysis to regions of the cell, where the spatially filtered cell centerline (using a Gauss filter of *σ* = 80 nm) was nearly straight (Fig. 7A). These regions still showed variations of cell-envelope curvature due to bulges or indentations (Fig. 7F). However, we did not find any significant correlations between MreB and contour curvature (Fig. 7D). Therefore, all residual MreB-curvature correlations after removal of poles and septa can be attributed to weak cell bending, while bending-independent bulges and indentations do not affect MreB localization.

We confirmed our findings with two alternative approaches: First, we subtracted from the local contour curvature the curvature contribution due to cell bending (Fig. 7–SI Fig. 2A-B). Residual curvature fluctuations originate from bulges or indentations. Consistently with our observation in straight cell segments, we found no significant correlations between MreB and corrected contour curvature. In an independent approach, we corrected MreB intensity values along the contours for the effect expected from cell-centerline bending (Fig. 7–SI Fig. 2C). Again, we did not find residual correlations after correction. Both of these analyses therefore strongly support the conclusion that spontaneous cell bending is responsible for all correlations between MreB and contour curvature in the cylindrical parts of normally growing cells.

Previously, we demonstrated that a small bending-induced enrichment of MreB can be explained by persistent rotational motion of MreB filaments (Wong et al., 2017), because rotating MreB filaments tend to accumulate at inner regions of bent cells. Our observations are therefore compatible with a model of MreB-independent initiation of Rod complexes through PBP2.

### Diffusing PBP2 molecules do not contribute to Rod-complex activity

It was previously suggested that diffusive PBP2 molecules contribute to cell-wall synthesis (Lee et al., 2014). However, if diffusive PBP2 molecules indeed contributed to processive Rod-complex activity, any cross-linking site of a moving Rod complex would have to be found by independent PBP2 molecules through diffusion at a rate equal to the cross-linking rate of up to 15/s. This rate corresponds to the distance between cross links of 2 nm (Meroueh et al., 2006) and a speed of PBP2 of 30 nm/s frequently observed (Fig. 1F).

To test whether PBP2 diffusion was fast enough for the biological cross-linking rate, we conducted computational simulations of freely diffusing PBP2 molecules and measured the average encounter rate between any simulated molecule and a given target site representing a Rod complex (Fig. 8A, B). We found that the rate between encounters was at least three times lower than the rate of cross-linking, even if a single enzyme had a high chance of returning to the same site and conduct multiple cross-linking reactions on average (Fig. 8C). Therefore, free diffusion cannot account for physiological cross-linking rates.

**Figure 8.**
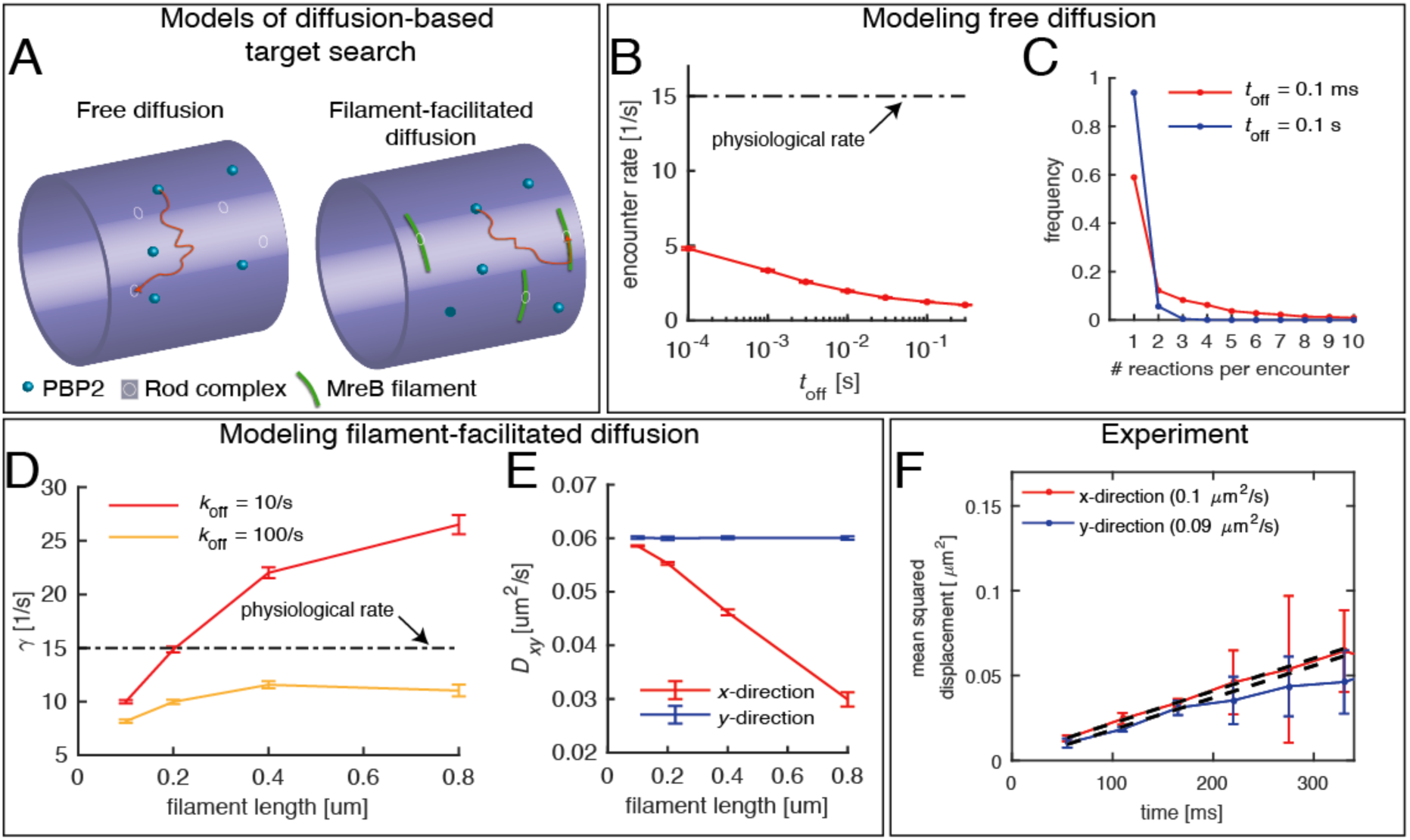
Testing a possible role of diffusive PBP2 for cross-linking. **(A)** Cartoon of PBP2 finding the target site of a ‘rod complex’ through free diffusion (left) or filament-facilitated diffusion (right). **(B)** The average encounter rate between any of 100 freely diffusing PBP2 molecules and a given rod-complex site as a function of the unknown latency time *t*_off_ (the duration for which a single PBP2 enzyme is inactive after a cross-linking reaction) (red) in comparison to the physiological cross-linking rate (dashed-dotted line) **(C)** Distribution of the number of successive cross-linking reactions conducted by the same PBP2 molecule at the same rod-complex site for two different latency times. **(D-E)** Facilitated diffusion along circumferentially oriented filaments centered at every rod-complex site increases the encounter rate **(D)** and renders diffusion asymmetric **(E)**. **(F)** Diffusion of PAmCherry-PBP2 is not asymmetric at short time scales, suggesting that PBP2 does not undergo facilitated diffusion along filaments.

As an extension of the model we considered the possibility that PBP2 molecules underwent facilitated diffusion by preferentially diffusing along MreB filaments, which could possibly serve to increase encounter rates between PBP2 and cell-wall-insertion sites. One-dimensional diffusion along filaments indeed increased the encounter rate (Fig. 8D). However, due to the preferentially circumferential orientation of MreB filaments, facilitated diffusion would also lead to reduced and asymmetric diffusion (Fig. 8E). On the contrary, we did not observe asymmetric diffusion in our experiments (Fig. 8F). Therefore, Rod-complex activity requires the stable association between transglycosylase and transpeptidase for multiple cross-linking events. Our findings suggest, that only the persistently moving fraction of PBP2 molecules substantially contributes to cross-linking.

## Discussion

In summary, we found a dominant role of PBP2 for Rod-complex initiation and persistent cell-wall synthetic activity. New Rod complexes are initiated by PBP2 binding to an immobile substrate that is different from any known Rod-complex component and likely the cell wall. While MreB filaments are required for Rod-complex motion, they have no influence on the initial location of PBP2 binding. Therefore, PBP2 determines the location of newly forming Rod complexes independently of MreB and likely independently of all other known Rod-complex components.

It has been proposed that Rod complexes are recruited to sites of specific cell-envelope curvature based on mechanical properties of MreB (Ursell et al., 2014). We found that correlations between MreB filaments and cell-envelope curvature in normally growing rods do not require any curvature-dependent initiation of Rod complexes. This observation does not rule out that MreB-curvature correlations in cells of strongly perturbed shape might be influenced by MreB-filament bending or twisting (Bratton et al., 2018; Colavin et al., 2018). Evidence for motion-independent curvature preferences comes from *C. crescentus* (Harris et al., 2014). However, our study as well as previous studies (Hussain et al., 2018; Wong et al., 2017, 2019) suggest that MreB-filament rotation around the circumference are responsible for MreB-curvature correlations in wild-type and filamentous cells. Therefore, the physical signal underlying the spatial pattern of new Rod complexes is likely found in the local architecture of the cell wall, and not, as previously suggested, in the geometry of the cytoplasmic membrane.

We found that the bound fraction of PBP2 molecules remained nearly constant upon A22 or Mecillinam treatment (Fig. 3). We thus reasoned that persistently moving and immobile molecules are likely bound to the same substrate. In Gram-negative *E. coli* active Rod complexes are thought to insert nascent glycan strands in between template strands (Höltje, 1998), even if deviations from perfect alignment are reported (Turner et al., 2018). During cell-wall insertion, Rod complexes might therefore stay connected to the local cell wall through associations between PBP2 and a template strand, independently of enzymatic activity. In the future, it will be interesting to study potential interactions at the molecular level. These might then also reveal structural features of the cell wall potentially responsible for stable PBP2 association.

PBP2 molecules transition only slowly between bound and diffusive states (Fig. 2). Possibly, PBP2 is found in two different molecular states that facilitate stable binding or allow for diffuse motion – either through conformational change or through interaction with an unknown interaction partner. Here, we demonstrated that depletion of RodA or MreC and MreD did not reduce the bound fraction of bound PBP2 molecules despite pronounced effects on cell shape, suggesting that RodA, MreC, and MreD have no major role on PBP2 binding or on stabilizing bound PBP2. RodZ depletion, on the contrary, showed a reduction of the bound fraction, suggesting that RodZ directly modulates PBP2 binding or the stability of the bound form of PBP2. This is compatible with previous observations of RodZ-PBP2 interactions (Bendezú et al., 2009; Morgenstein et al., 2015). In the future, it will be interesting to investigate the role of RodZ for PBP2 binding or unbinding in more detail.

How do MreB filaments ‘find’ bound PBP2 molecules? MreB filaments are generally thought to move slowly. However, small filaments might also undergo rapid diffusive motion previously overlooked. Alternatively, PBP2 – either alone or together with other Rod-complex components such as RodZ – could nucleate new MreB filaments. This latter hypothesis is supported by the recent observation that the hyperactive PBP2(L61R) mutant described above causes more and shorter MreB filaments (Rohs et al., 2018).

We found that PBP2 positively limits processive Rod-complex activity, as demonstrated by nearly constant fractions of bound and persistently moving PBP2 molecules upon variations of PBP2 levels, and by high residual numbers of persistently moving PBP2 upon depletion of RodZ, MreCD, or RodA. These observations challenge the idea of a well-defined complex that requires the presence of all components for processive cell-wall-synthetic activity. RodA has been proposed as the major transglycosylase of the Rod complex (Emami et al., 2017; Meeske et al., 2016). Based on (Vigouroux et al., 2018), RodA levels were reduced more than 6-fold with respect to native levels in our experiments, while PAmCherry-PBP2 was overexpressed by two- to four-fold (Fig. 1–SI Fig. 1D). Yet, the number of persistently moving molecules per cell (bound fraction times persistent fraction) dropped by less than two-fold between wildtype and RodA repression. With a wild-type stoichiometry between RodA and PBP2 of about 1.4 (Li et al., 2014) there should thus have been less than one RodA molecule per moving PBP2 molecule. The high residual degree of persistent motion thus suggests that a different transglycosylase can be part of the active Rod complex, either a class-A PBP such as PBP1a (Banzhaf et al., 2012) or the SEDS protein FtsW (Taguchi et al., 2019).

Despite the substantial residual degree of persistent motion upon depletion of MreCD, RodA, or RodZ, the presence of all these components is required to stably maintain rod shape. A recent paper suggested that cell diameter in *E. coli* is determined by the spatial density of active Rod complexes (Dion et al., 2019). The model is compatible with the fold-change of persistently moving PBP2 molecules and cell-diameter changes observed during MreCD and RodZ depletion. However, during RodA-depletion this simple model does not apply: Due to overexpression of PBP2, the total number of persistently moving PBP2 molecules per cell is as high as in the wildtype – even during RodA depletion. In the future, it will therefore be interesting to study how the stoichiometry of different Rod-complex components affects cell-wall insertion and cell shape.

## Acknowledgements

We thank Timothy Lee and KC Huang for strain TKL130 and plasmid pKC128, Nikolay Ouzounov and Zemer Gitai for strains NO50 and NO53, Tom Bernhardt for strains TU230(attLHC943) and TU230(attLPR122), Felipe Bendezú and Piet de Boer for plasmids pFB121, pFB290 and strain FB60(iFB273). This work was supported by the European Research Council (ERC) under the Europe Union’s Horizon 2020 research and innovation program [Grant Agreement No. (679980)], the French Government’s Investissement d’Avenir program Laboratoire d’Excellence “Integrative Biology of Emerging Infectious Diseases” (ANR-10-LABX-62-IBEID), the Marie de Paris “Emergence(s)” program, the PRESTIGE Postdoc fellowship (Campus France), and the Volkswagen Foundation.

