## Supplemental Figures for "Transpeptidase PBP2 governs initial localization and activity of major cell-wall synthesis machinery in *Escherichia coli*"

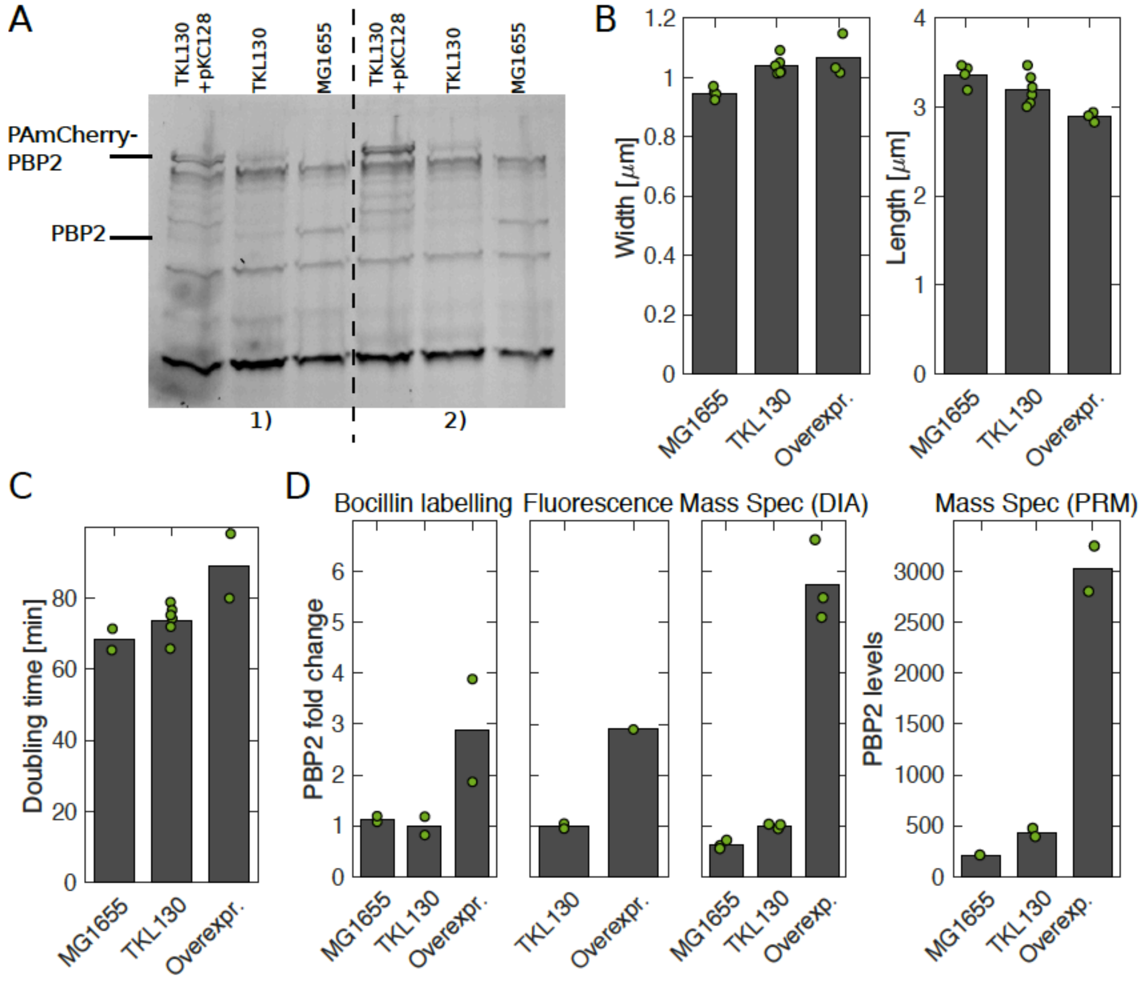


**Figure 1–figure supplement 1. Comparison of PBP2-PAmCherry expressing cells and WT.**

**(A)** Bocillin-binding assay to compare expression levels of PBP2 in the wild-type strain (MG1655), the strain expressing PBP2-PAmCherry from the native locus (TKL130), and the strain overexpressing PBP2-PAmCherry (TKL130/pKC128). Two replicates. Quantification in (D) **(B)** Average cell dimensions obtained by phase-contrast microscopy and computational image segmentation. **(C)** Average doubling times during steady-state exponential growth in batch culture (from OD600). **(D)** Different methods to compare PBP2 expression levels in different strains (from left to right): Bocillin labeling (from A), single-cell fluorescence levels measured in epi-fluorescence mode, mass spectrometry [Data Independent Acquisitions (DIA) and Parallel Reaction Monitoring (PRM)]. For the first three methods, PBP2 levels are normalized by the corresponding value in TKL130. For PRM, we obtained absolute numbers of proteins per cell by comparing to reference peptides and colony counting. With both mass spectrometry methods, we observe a higher fold-change than through the other methods. Dots represent independent replicates.

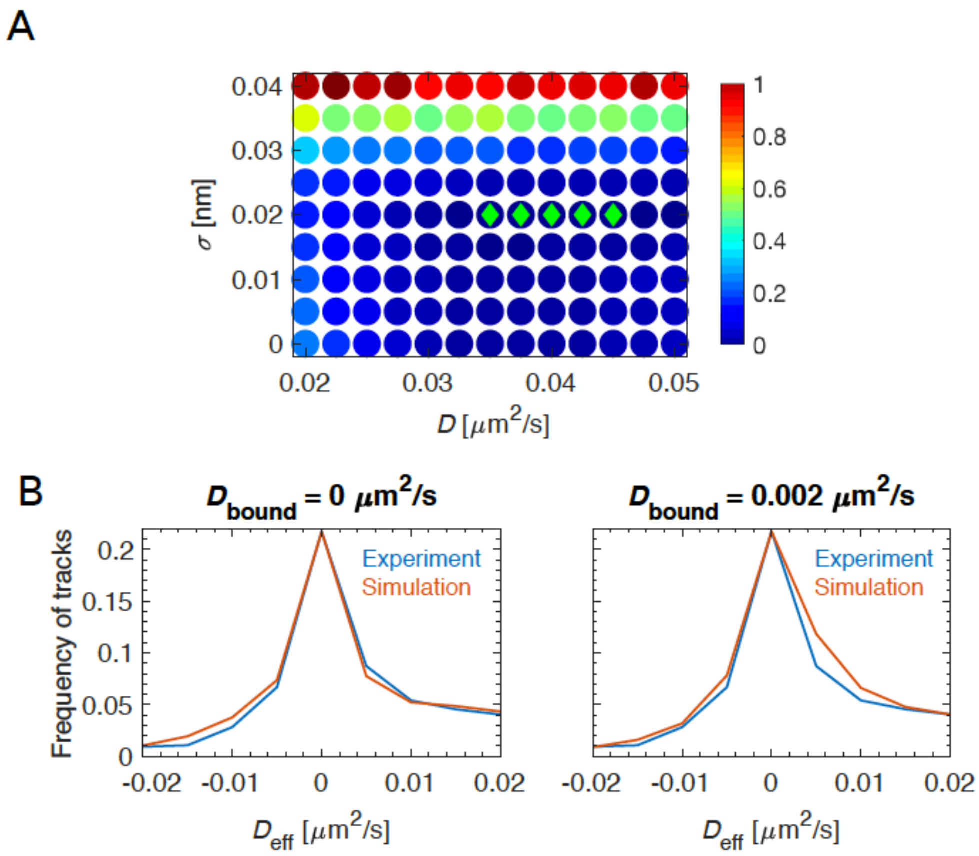


**Figure 1–figure supplement 2. Fitting a two-state diffusion model.**

**(A)** Heat map of residual sum of squared differences (RSS) between $D_{eff}$ distributions of simulation and experimental data for different parameters $D$ and $\sigma$. Parameter sets giving the lowest 5 RSS values are shown with green diamonds. Best fit is given by $D$ = 0.04 μm$^2$/s and $\sigma$ = 20 nm. **(B)** We verified that the non-diffusive population was indeed not diffusing, with $D_{bound}$ = 0 μm$^2$/s (left), while a finite diffusion constant $D_{bound}$ > 0.002 um$^2$/s gives poor agreement between simulation and experiment. Here, the experimental $D_{eff}$ distribution is the mean of 6 independent replicates.

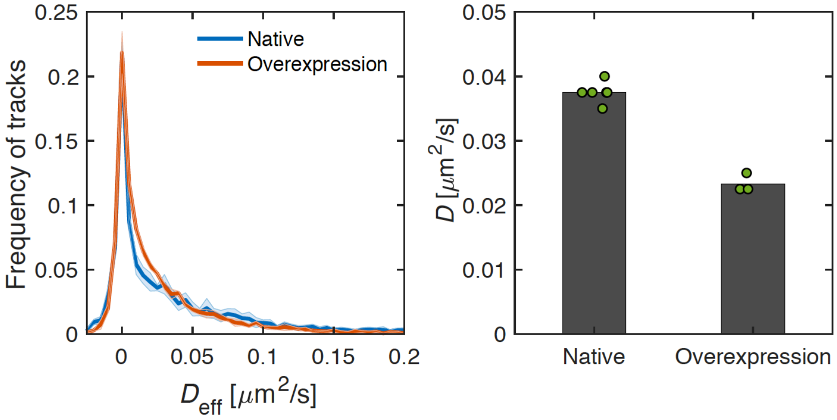


**Figure 1–figure supplement 3. Overexpression of PAmCherry-PBP2.**

Distributions of effective diffusion coefficients (left) and extracted diffusion constants of the diffusive population (right) for PAmCherry-PBP2 expressed at native levels in TKL130 or overexpressed in TKL130/ pKC128. Shaded region shows standard deviation between replicates.

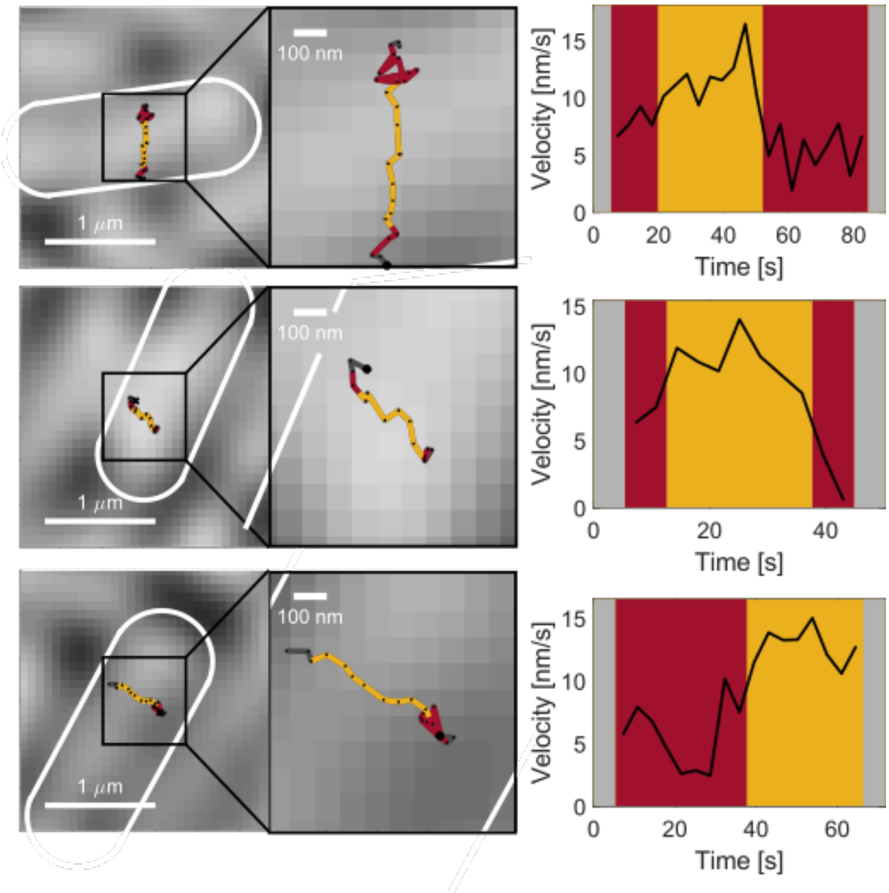


**Figure 1–figure supplement 4. Transitions between immobile and persistent states.**

Examples tracks and velocity as a function of time for example tracks that show transitions between persistent and immobile states.

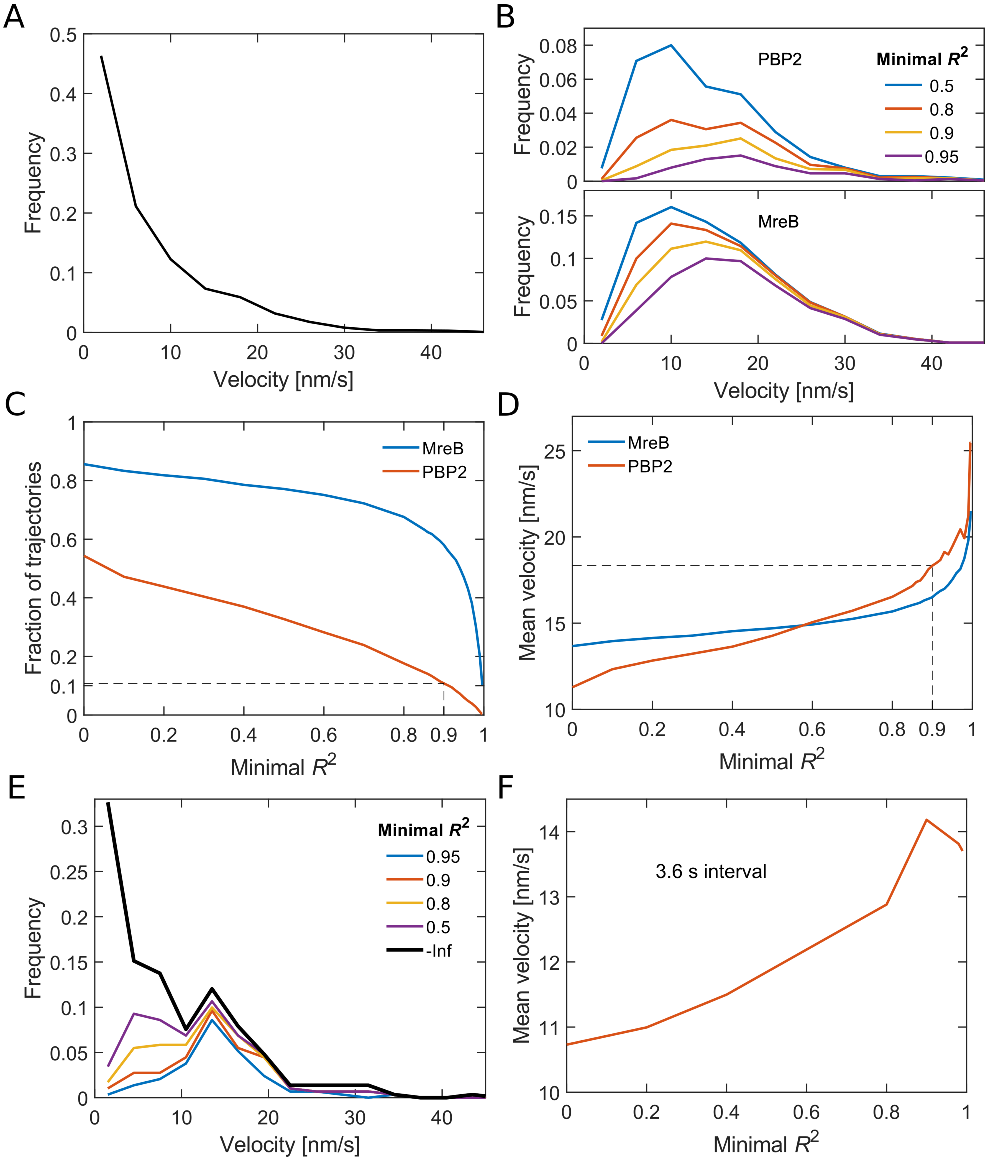


**Figure 1–figure supplement 5. Analysis of bound PAmCherry-PBP2 molecules.**

**(A)** Velocity distribution of all PBP2 tracks measured with 1 s intervals. The velocity of individual tracks was determined by fitting a quadratic function to the MSD. **(B)** Velocity distributions for directed trajectories of PBP2 and MreB as found by selecting for an increased goodness of fit measured by $R^2$ of a quadratic function to the MSD. **(C)** The stricter the goodness of fit criterion (minimal $R^2$) the less trajectories contribute to the mean track velocity. **(D)** The mean velocity increases with increasing minimum $R^2$. The dashed line indicates the value chose for the distributions in Fig. 1. **(E-F)** The same analysis applied on 4-step segments of trajectories measured with 3.6 s intervals delivers smaller mean velocities, likely because fast trajectories reside for a shorter amount of time in the field of view.

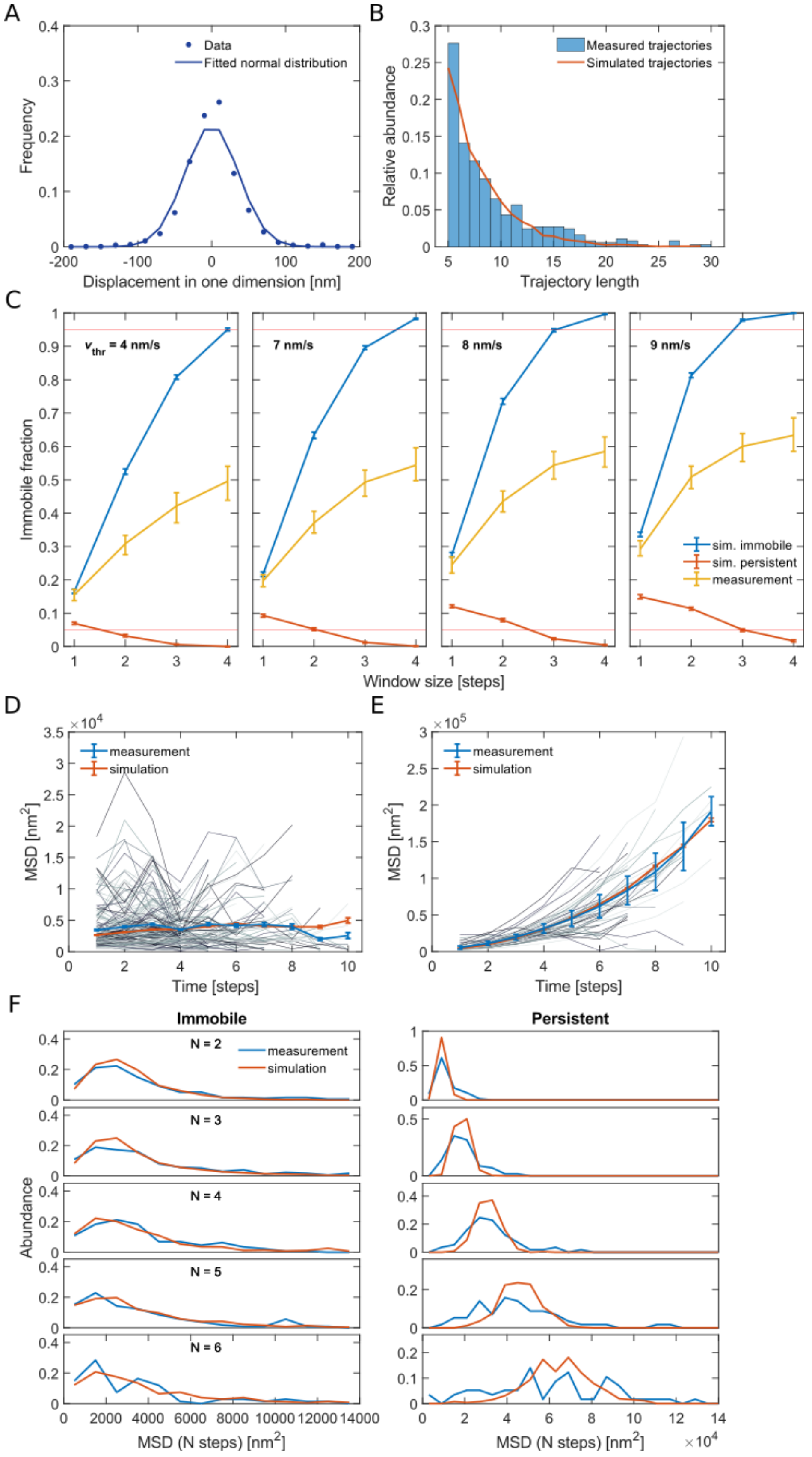


**Figure 1–figure supplement 6. Quantitative analysis of persistent and immobile states based on computational simulations.**

**(A)** Distribution of measured single-step displacements in one dimension. A fit of a normal distribution to the data delivers a standard deviation of 36 nm, which corresponds to a localization error of single localization events of 25 nm. **(B)** We computationally simulated trajectories such that the length distribution of the simulated trajectories resembles the one from measured trajectories. **(C)** Fraction of immobile segments measured in simulations of immobile (blue) or persistent (red) molecules and in experimentally measured tracks (yellow) as a function of the moving-average window size and for different velocity thresholds. The red horizontal lines signify 5% and 95% probability thresholds, respectively. Error bars are from bootstrapping. For a window size of 4 steps and a velocity threshold of 8 nm/s the rate of wrong annotation is smaller than 1% both in simulations of purely persistent or immobile molecules. For pairs of $w$ and $v_{thr}$ that lead to high accuracy of the determination of immobile and persistent segments the immobile fraction of the experimental data shows similar results. **(D-E)** MSD's of single-track segments (gray lines) classified as **(D)** immobile or **(E)** persistent compared to the MSD of all respective segments (blue line). For simulated trajectories that can switch between the immobile and the persistent state (simulated with $v$ = 12 nm/s, $k_{ip}$ = 0.015 $s^{-1}$, $k_{pi}$ = 0.021 $s^{-1}$) we find a similar behavior of the MSD curves (red line). **(F)** Distribution of MSD's of immobile and persistent segments for different numbers of steps $N$.

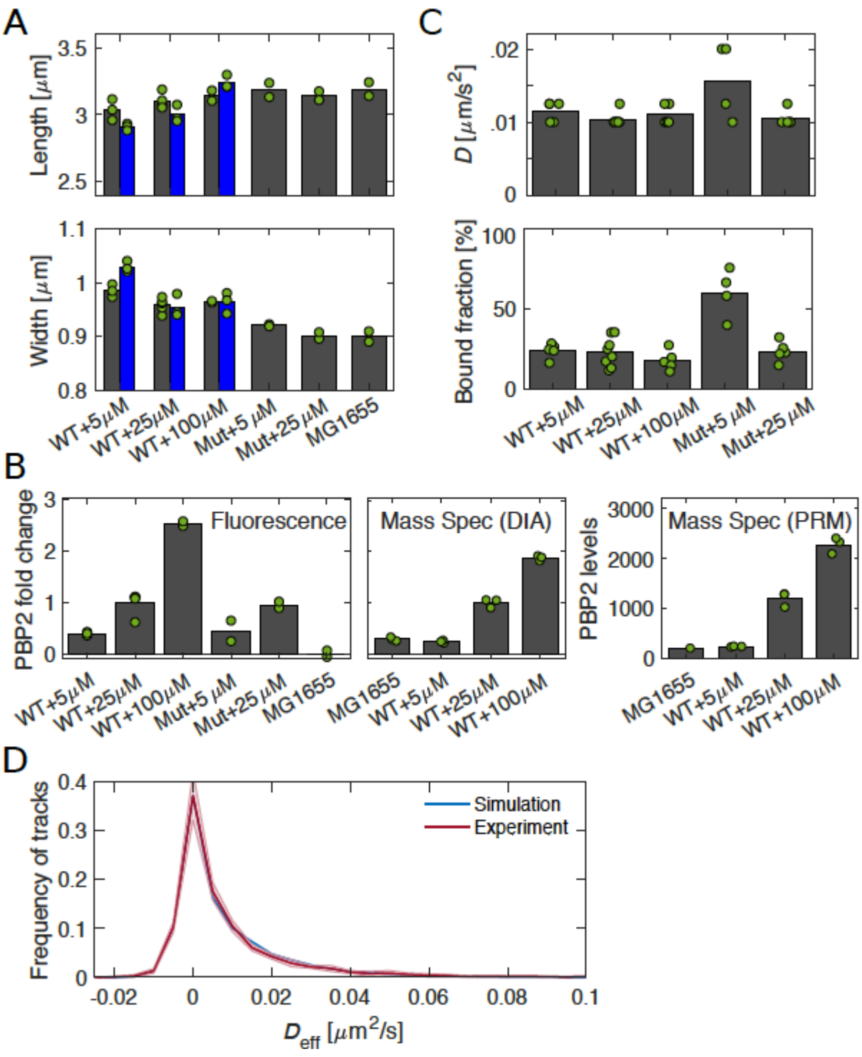


**Figure 1–figure supplement 7. Comparison of msfGFP-PBP2 (TU230(attLHC943)), msfGFP-PBP2(L61R) (TU230(attLHC943)), and WT strains.**

**(A)** Width and length of msfGFP-PBP2 (labeled 'WT') and msfGFP-PBP2(L61R) (labeled 'Mut') for different induction levels in comparison to MG1655. Gray and blue bars show cell dimensions after 6 and 10 hours of growth, respectively (see also Fig. 1–SI Fig. 8). **(B)** Corresponding average fluorescence intensities for conditions in (A) from epi-fluorescence measurements and mass spectrometry measurements (DIA and PRM). PRM measurements combined with colony counting yield absolute numbers of proteins per cell. **(C)** Diffusion constants and bound fractions. Gray bars show data after 6 hours of growth. Dots represent independent experiments. **(D)** Distributions of effective diffusion coefficients for simulated and experimental data. Shaded regions represent standard deviation between replicates.

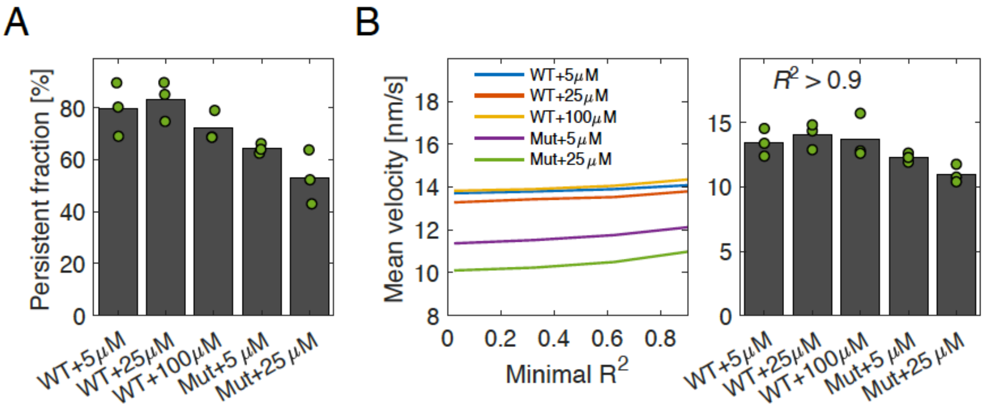


**Figure 1–figure supplement 8. Low-frequency tracking of msfGFP-PBP2 and msfGFP-PBP2(L61R) cells under different induction levels.**

**(A)** Persistent fractions **(B) Left.** Mean velocity as a function of minimal $R^2$ which are obtained from a quadratic fit to MSD of the form $y = a+bx^2$ **Right.** Mean velocity of tracks which satisfy $R^2>0.9$ are shown. Dots represent independent experiments.

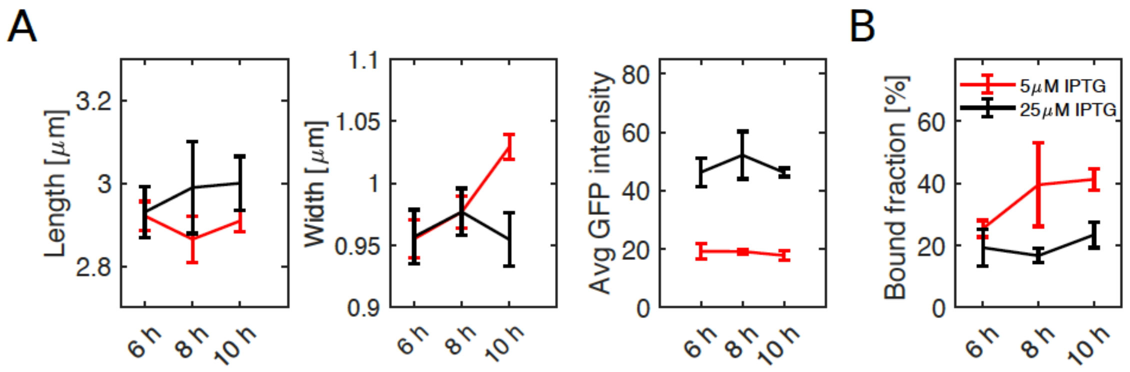


**Figure 1–figure supplement 9. Change of PBP2 levels as a function of time.**

**(A)** Cell length, width and GFP intensity and **(B)** bound fractions of TU230(attLHC943) at 6, 8 and 10 hours of growth for 2 different induction levels of 5 and 25 μM IPTG. Error bars show standard deviation between independent experiments.

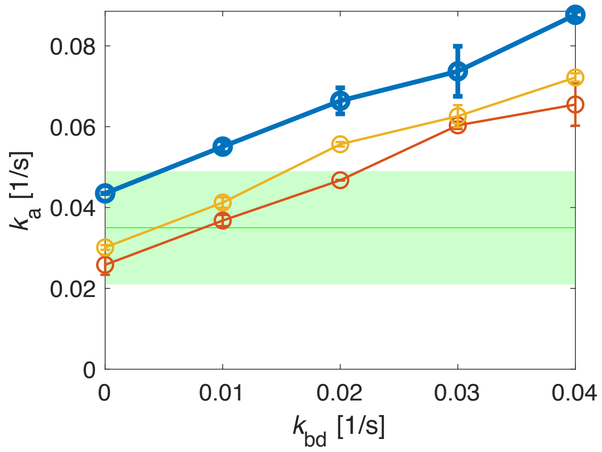


**Figure 2–figure supplement 1. Determination of an upper limit of the unbinding rate $k_{bd}$ through simulations.**

Simulations of track-length distributions reveal the apparent track termination rate $k_a$ as a function of the unbinding rate $k_{bd}$ for different transition simulated rates $k_{ip}$, $k_{pi}$. Top: $k_{ip}$ = 0.063/s, $k_{pi}$ = 0.0086/s (experimentally measured rates, leading to a bound fraction of 88%); middle and bottom (thin solid lines): $k_{ip}$ = 0.063/s, $k_{pi}$ = 0.0158/s; $k_{ip}$ = 0.033/s, $k_{pi}$ = 0.0086/s. For the thin solid lines, we adjusted either of the two rates to yield the experimentally measured bound fraction of 80%. Comparison with the experimentally measured apparent unbinding rate (green) allows us to infer an upper bound for the unbinding rate $k_{bd}$ < 0.03/s. Shaded area: 95% confidence interval.

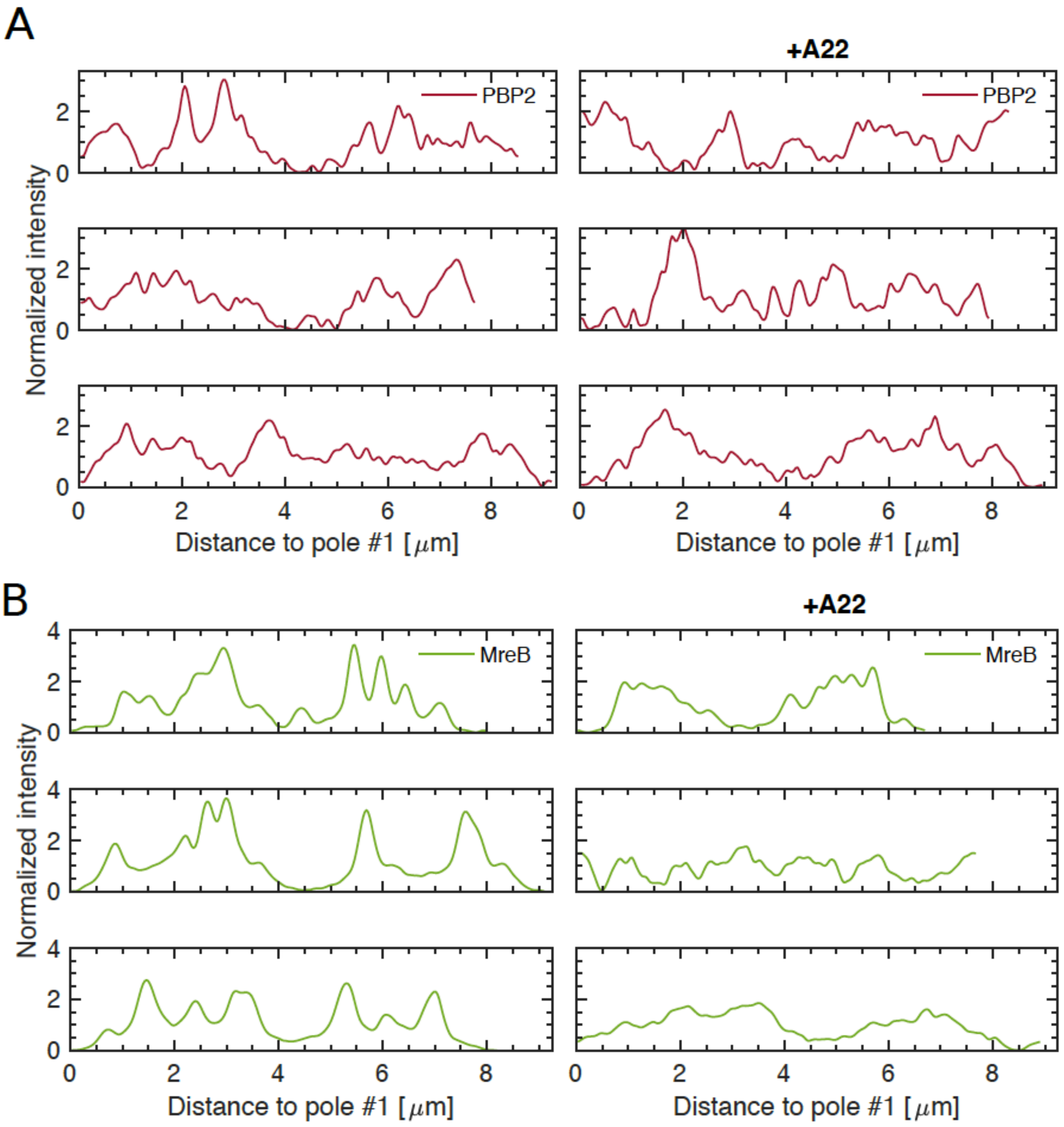


**Figure 3–figure supplement 1. A22 treatment does not affect PBP2 localization.**

Fluorescence profiles along contours of different cells carrying mCherry-PBP2 **(A)** or MreB-msfGFP **(B)** fusions for untreated **(left)** or A22 treated cells (50 μg/ml) **(right)**, obtained in the same way as in Fig. 3A, B. Intensities are normalized by the median value and smoothened with a Gauss filter with standard deviation of 33 nm (0.5 pixel).

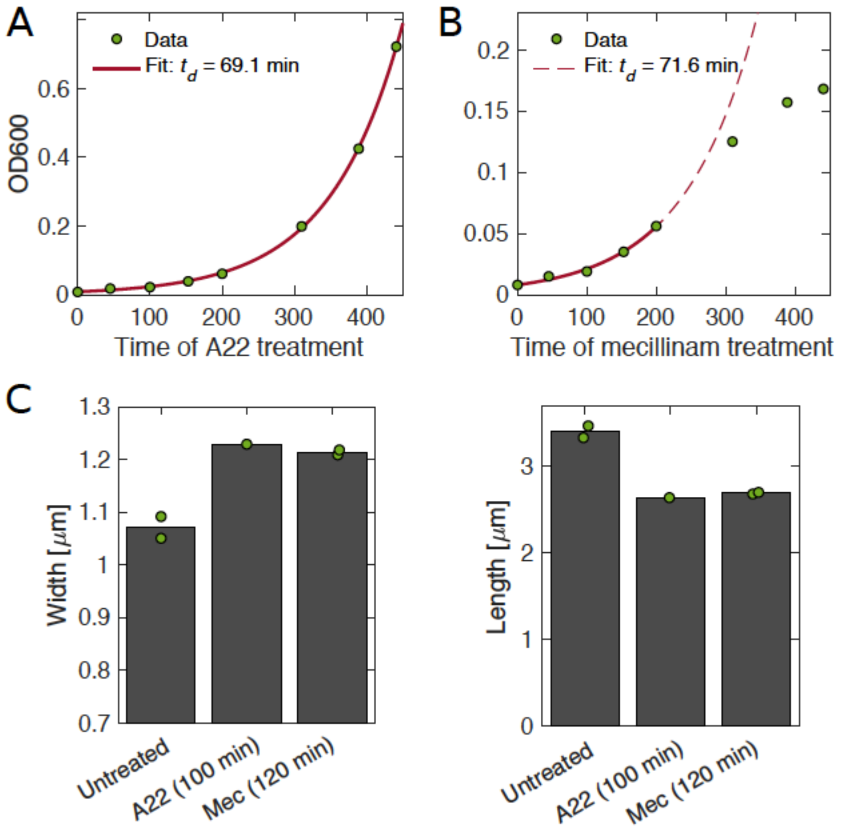


**Figure 3–figure supplement 2. Growth and shape of A22- and mecillinam-treated cells.**

**(A-B)** A22-treated cells grow unperturbed for 6 generations (A), while cells treated with mecillinam show a reduced growth rates after around 3 generations (B). **(C)** Cell shape of cells treated with A22 (C) and mecillinam (B). In both cases**,** cells become wider and shorter. Dots represent independent experiments.

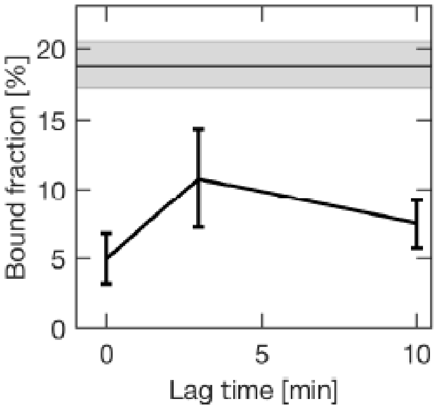


**Figure 4–figure supplement 1. Change in the bound fraction after photobleaching of untreated cells.**

The horizontal line corresponds to the mean bound fractions obtained from unbleached cells (see Fig 3). Error bars and shaded area show standard deviations between replicates.

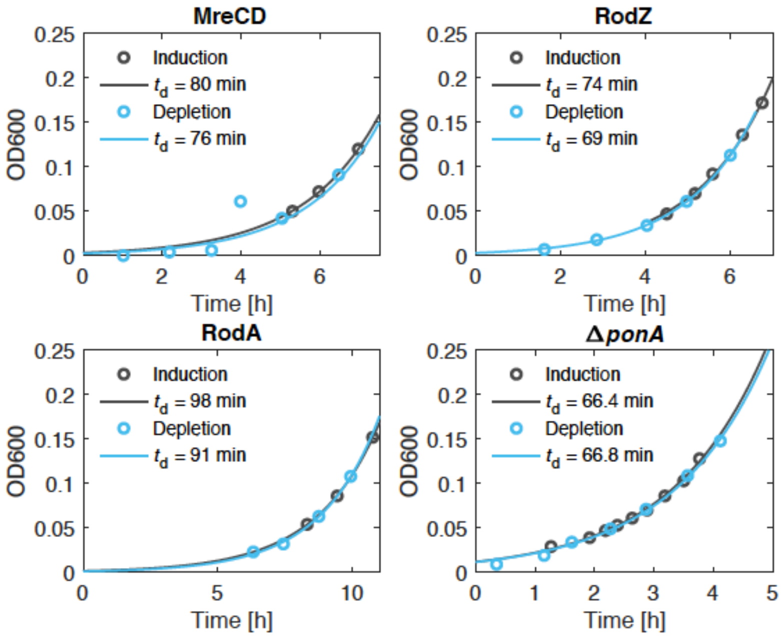


**Figure 5–figure supplement 1. Depletion of Rod-complex components shows no effect on growth.**

Growth curves for the different depletion strains in induced and deplete conditions as a function of time after initiating protein depletion. The doubling time is obtained from an exponential fit. For the fit to the data of the MreCD depletion we did not consider the point at 4h.

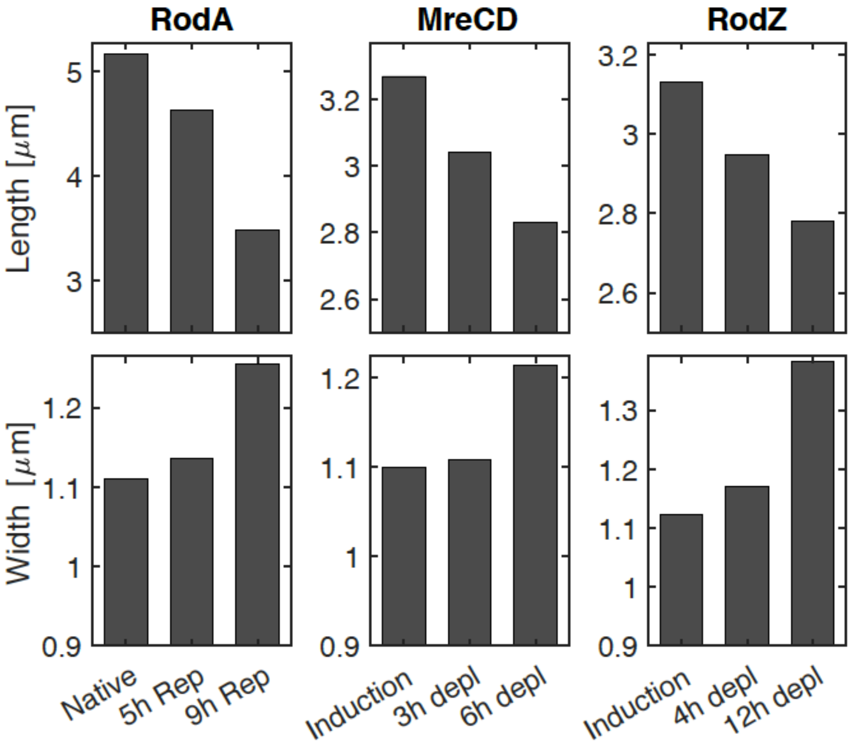


**Figure 5–figure supplement 2. Depletion of Rod-complex components leads to loss of rod shape.**

Cell length (top) and width (bottom) upon repression of *rodA* (left), and depletion of MreCD (middle) or RodZ (right) at different time points after initiating protein depletion.

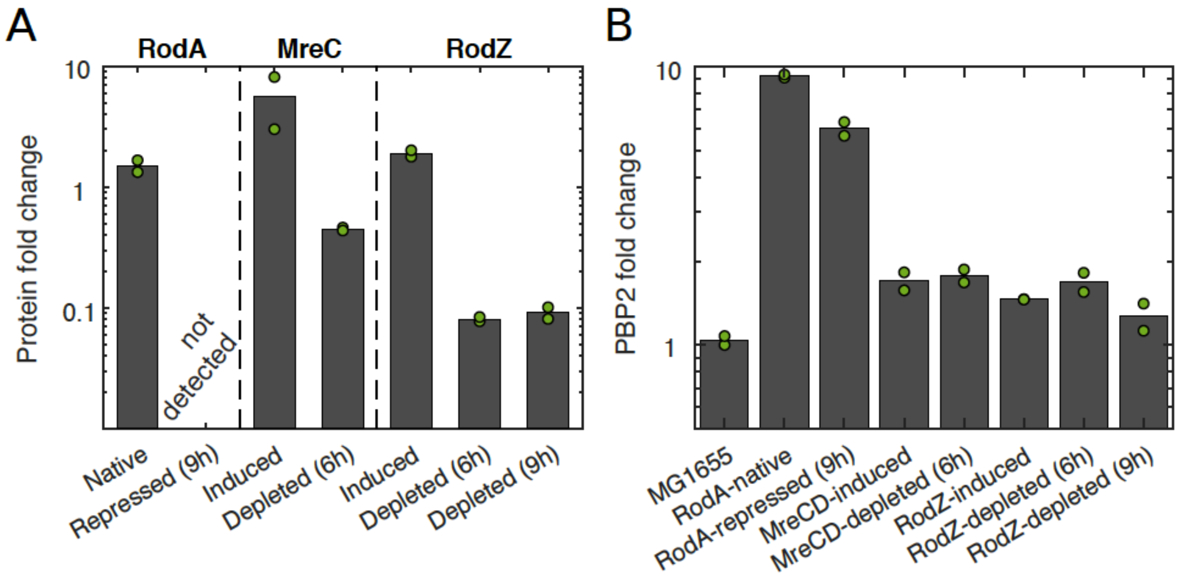


**Figure 5–figure supplement 3. Protein levels during depletion of Rod-complex components.**

**(A-B)** Levels of RodA, MreC, and RodZ (A) and levels of PBP2 (B) acquired by mass spectrometry (DIA). Protein levels are normalized by the mean of the corresponding protein level in MG1655. Two biological replicates for each condition. Dots represent independent experiments.

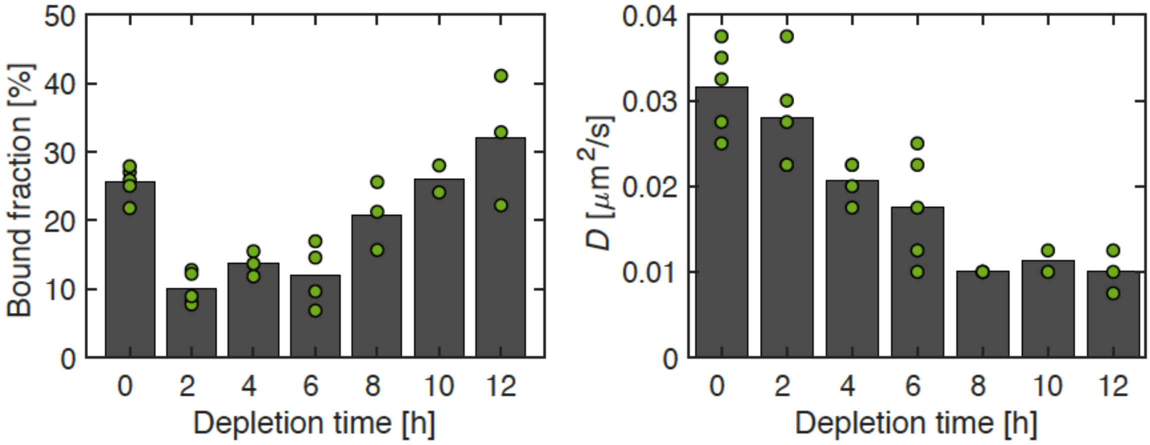


**Figure 5–figure supplement 4. Effect of RodZ depletion on PBP2 dynamics.**

Bound fraction and diffusion constant of PBP2-PAmCherry at different time points during RodZ depletion. Dots represent independent replicates.

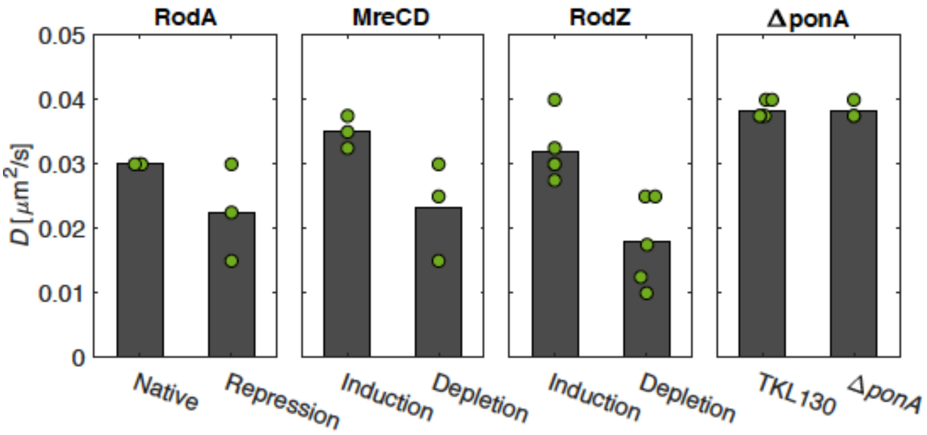


**Figure 5–figure supplement 5. Effect of the depletion of Rod-complex components on diffusion.**

Diffusion constants drop upon repression of potential members of the Rod complex. Dots represent independent replicates.

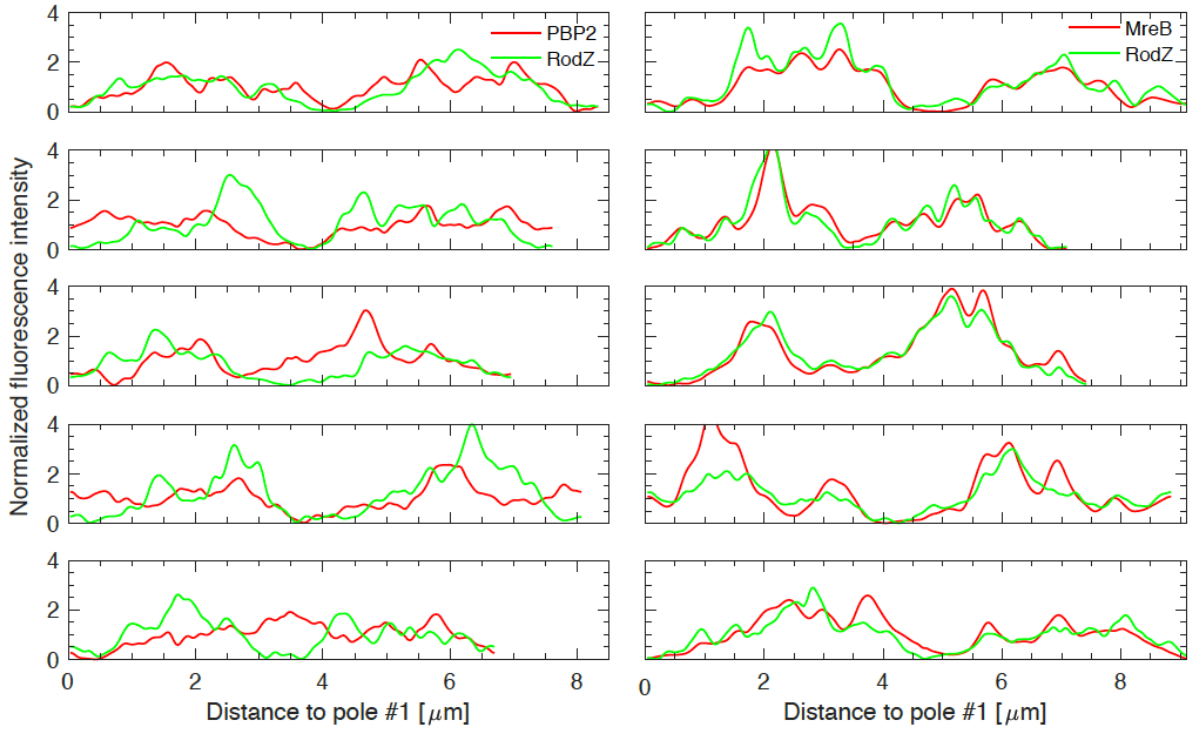


**Figure 6–figure supplement 1. RodZ and PBP2 do not colocalize, while RodZ and MreB do.**

Sample profiles of different cells carrying mCherry-PBP2 and GFP-RodZ fusions (Δ*rodZ mrdA*<>*mCherry-mrdA* ($P_{lac}$::*gfp-rodZ*)) **(left)** or MreB-mCherry and GFP-RodZ fusions (Δ*rodZ mreB*<>*mreB-mCherry* ($P_{lac}$::*gfp-rodZ*)) **(right)** as in Fig. 6C. Intensities are normalized by the median value and then smoothened with a Gauss filter with standard deviation of 33 nm (0.5 pixel).

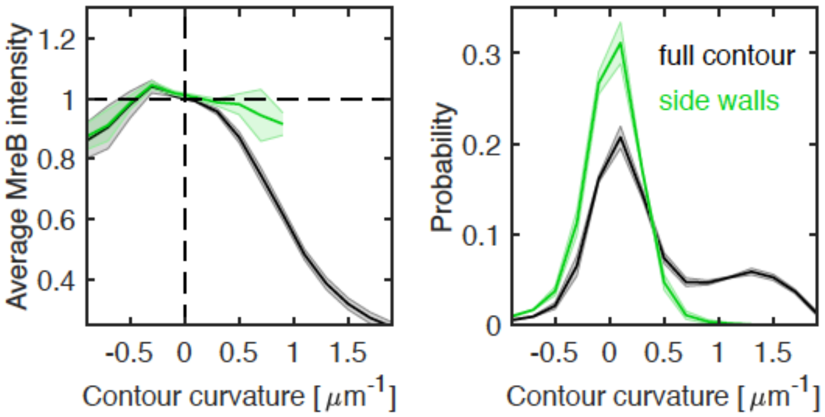


**Figure 7–figure supplement 1. Correlations between MreB and contour curvature in WT cells.**

**Left.** Normalized average MreB intensity as a function of local contour curvature in strain NO53 (*mreB<>mreB-msfGFP*). Comparison between correlations obtained from full contours (black) and side walls (green). **Right.** Distributions of contour-curvature values corresponding to correlation plots on the left.

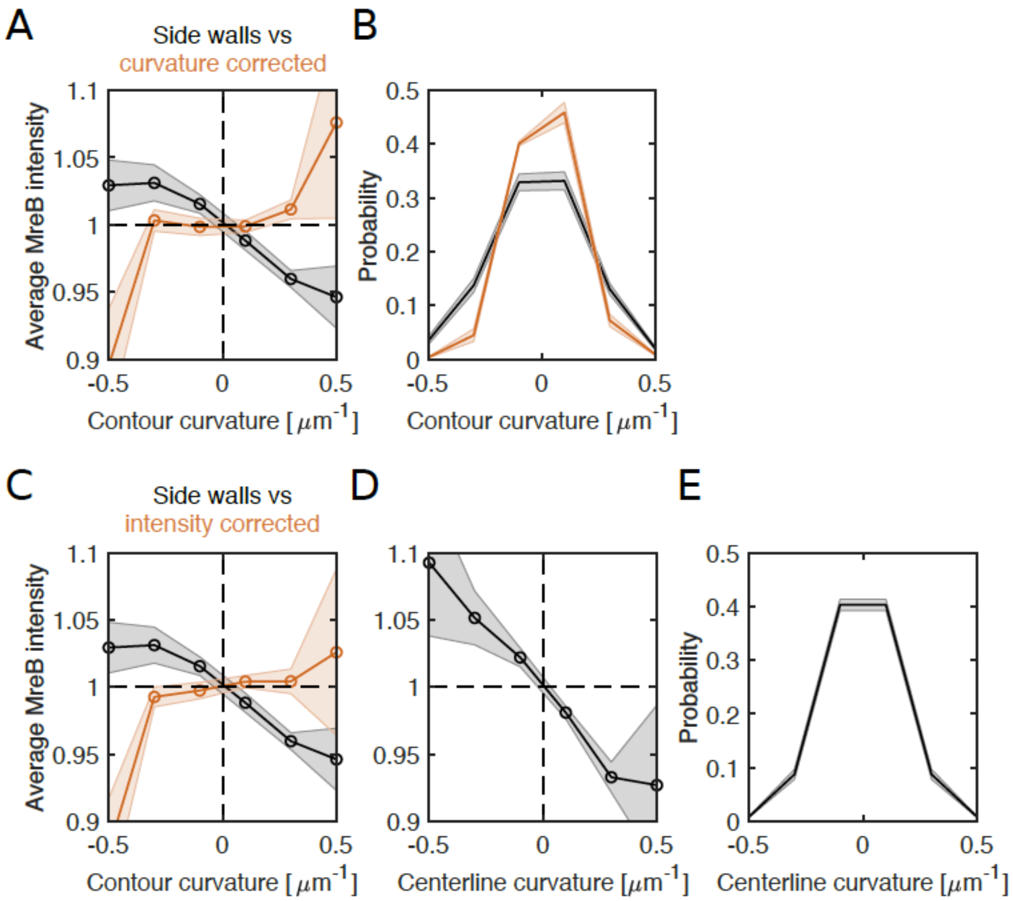


**Figure 7–figure supplement 2. Loss of correlations between MreB and contour curvature after renormalizing either curvature or intensity for cell bending in filamentous cells.**

**(A-B) Curvature correction. (A)** Average MreB intensity as a function of contour curvature (black) and bending-corrected contour curvature (orange) in NO53/pDB192 ($P_{lac}$::*sulA*). **(B)** Distributions of contour curvature (black) and corrected contour curvature (orange). Curvature correction leads to a narrower distribution while ~80% of local curvature values keep their original sign of curvature.

**(C-E) Intensity correction. (C)** Average MreB intensity as a function of bending-corrected MreB-intensity (orange). Intensity is corrected for observed correlations between MreB intensity and centerline curvature in (D) (Methods). **(D)** Average MreB intensity as a function of smoothened centerline curvature (using a Gauss filter of $\sigma$ = 80 nm). **(E)** Centerline-curvature distribution.

### Movie captions:

**Fig. 1–Movie 1. High-frequency imaging of PAmCherry-PBP2.**

**Left.** Denoised bright-field image taken at the beginning of the movie. **Right.** Raw images from high-frequency imaging (imaging interval 60 ms) of PAmCherry–PBP2 show diffusive and bound molecules. Blue circles and lines represent peaks and corresponding tracks considered for analysis.

**Fig. 1–Movie 2. Low-frequency imaging of PAmCherry-PBP2.**

**Left.** Denoised bright field image **Right.** Raw images from low frequency imaging (imaging interval 3.5 s) of PAmCherry–PBP2 show immobile and persistently moving molecules. Blue circles and lines represent peaks and corresponding tracks considered for analysis.

**Fig. 1–Movie 3. High-frequency imaging of msfGFP-PBP2.**

**Left.** Denoised bright field image taken at the beginning of the movie**. Right.** Raw images from high frequency imaging (imaging interval 60 ms) of msfGFP–PBP2 show diffusive and bound molecules. Blue circles and lines represent peaks and corresponding tracks considered for analysis.

**Fig. 1–Movie 4. Low-frequency imaging of msfGFP-PBP2**

**Left.** Denoised bright field image **Right.** Raw images from low frequency imaging (imaging interval 3.5 s) of msfGFP–PBP2 show immobile and persistently moving molecules. Blue circles and lines represent peaks and corresponding tracks considered for analysis.

**Fig. 3–Movie 1. MreB-msfGFP imaging**

**Left.** Phase contrast image taken at the beginning of the movie. **Right.** Raw images of MreB-msfGFP motion (imaging interval 1 s) of NO53 cells. Blue circles and lines represent peaks and corresponding tracks considered for analysis.

**Fig. 3–Movie 2. MreB-msfGFP imaging of A22 treated cells**

**Left.** Phase contrast image taken at the beginning of the movie. **Right.** Raw images of MreB-msfGFP motion (imaging interval 2 s) of NO53 cells treated with 20 µg/ml A22 for a duration of 30 minutes. Blue circles and lines represent peaks and corresponding tracks considered for analysis.
