## Supplementary material for "Transpeptidase PBP2 governs initial localization and activity of major cell-wall synthesis machinery in *Escherichia coli*": Methods

### 1. Strain construction

EW07 To obtain the MreCD depletion strain EW07 (TKL130 Δ*mreCD*, pFB121) we introduced pFB121 (Bendezú and Boer, 2008) into TKL130. We replaced the *mreCD* gene by a kanamycin resistance gene (amplified from pKD13 with primer DmreC_fw and DmreC_rv) using the λ-Red mediated recombineering system expressed from pTKred (Kuhlman and Cox, 2010). The resulting strain was then grown at 42°C to cure the plasmid pTKred. We verified the deletion of *mreCD* by PCR.

EW49: For the RodZ depletion strain EW49 (TKL130 Δ*rodZ*, pFB290) we first introduced pFB290 (Bendezú et al., 2009) into TKL130. In the resulting strain we deleted *rodZ* by a P1 transduction with lysate made from the Δ*yfgA* strain from the Keio collection yielding strain EW49.

AV48: For RodA depletion we constructed a strain AV48 (186::$P_{tet}$-*dcas9*, *mrdA*::r*PAmcherry-mrdA*) that allows repressing the native mrdAB operon using CRISPRi and a guide RNA targeting the codon-modified ORF of rPAmcherry (Vigouroux et al., 2018), where 'r' denotes the codon modification (represible). The strain also carries the plasmid pKC128 ($P_{mrdA}$-*PAmcherry-mrdA*, non-repressible version) to counter repression of native PBP2. To construct strain AV48 we used allelic exchange on the strain LC69 (186attB::$P_{tet}$-*dcas9*) (Cui et al., 2018). To that end, we first electroporated the plasmid pAV25 (Vigouroux et al., 2018) into LC69 and selected for growth on chloramphenicol. Then we removed the plasmid backbone by transforming the cells with pAV10 (Vigouroux et al., 2018). The resulting strain was then grown at 42°C to cure the plasmid pAV10. We introduced the CRISPR plasmid pcrRNA G20-R20 (Vigouroux et al., 2018) and pKC128 into AV48.

S352: For the *ponA* deletion strain we deleted *ponA* in TKL130 by a P1 transduction with lysate made from the Δ*ponA* strain from the Keio collection yielding strain S352.

S504: To construct a strain expressing both MreB-mCherry and GFP-RodZ fusions, we used FB83 (*mreB-mCherry*) and performed P1 transduction with lysate made from FB60(iFB273) (Δ*rodZ*, *Plac::gfp-rodZ*) (Bendezú et al., 2009) first selecting on ampicillin for Plac::*gfp-rodZ* then a second transduction with the same lysate and selecting on kanamycin for deleting the native *rodZ*.

S505: To construct a strain carrying both mCherry-PBP2 and GFP-RodZ fusions, we used strain AV07 (*mCherry-mrdA*) and performed P1 transduction with lysate made from FB60 (iFB273) (Δ*rodZ*, *Plac::gfp-rodZ*) (Bendezú et al., 2009) first selecting on ampicillin for Plac::*gfp-rodZ* then a second transduction with the same phage lysate and selecting on kanamycin for deleting native *rodZ*.

### 2. Bocillin labelling

PBP2 levels of strains MG1655, TKL130 and TKL130/pKC128 were measured by a Bocillin-binding assay as described similarly in (Cho et al., 2016; Kocaoglu et al., 2012). We performed the quantification in parallel on two identical cultures grown from independent colonies of each strain. We grew strains overnight in LB at 37° C. We then washed the cells in LB, diluted them 1/200 in LB and grew them at 37° C until an OD600 of ~0.4. We washed 1.8 ml of the culture in PBS, resuspended in 200 µl PBS and kept cultures on ice. We disrupted cells by sonication (FB120, Fisher Scientific) and centrifuged them for 15 min at 4° C (21,000 g). We subsequently resuspended the pellet corresponding to the membrane fraction in 50 µl PBS containing 15 µM fluorescently labelled Bocillin-FL (Invitrogen). Membranes were incubated at 37° C for 30 min and washed once in 1ml PBS. We centrifuged the membranes for 15 min (21,000 g) and resuspended them in 50 µl PBS. We measured the protein concentration of each sample with a colorimetric assay based on the Bradford method (#5000006, Bio-Rad) and loaded equal amounts of protein mixed with 4X Laemmli buffer onto a 10 % polyacrylamide gel. We visualized the labelled proteins with a Typhoon 9000 fluorescence imager (GE Healthcare): excitation at 488 nm and emission at 530 nm. We quantified the relative amounts of PBP2 in each sample by quantifying the grey values of each lane in ImageJ (Schneider et al., 2012).

### 3. Sample preparation and imaging conditions

Cells were grown overnight at 37°C in LB medium and then washed and diluted at least 1:500 in M63 minimal medium (Miller, 1972) supplemented with 0.1% casamino acids, thiamine ($5 \cdot 10^{-5}$%), glucose (0.2%) and MgSO4 (1 mM) (here referred to as minimal medium) and grown to early exponential phase (maximum OD600 of 0.1) at 30°C. If strains carried antibiotic resistances we added the respective antibiotics to the medium (carbenicillin (100 µg/ml), kanamycin (50 µg/ml), chloramphenicol (30 µg/ml), all Sigma-Aldrich). Inducers were added as indicated below. For microscopy, cells were spotted on 1% agarose pads (UltraPure™ Agarose, Invitrogen) of minimal medium, without any antibiotics. Drugs targeting MreB or cell-wall synthesis and inducers were added as indicated below.

Cells were imaged in a custom-built temperature controlled chamber at 29°C. Prior to imaging the cells were incubated 10-20 min on the microscope to equilibrate sample temperature and minimize drift. In experiments with drugs [mecillinam (Sigma-Aldrich, #33447), A22 (Cayman Chemical #15870)] cells were spotted on agarose pads containing the antibiotic at the concentration of 100 µg/ml (mecillinam), or 20-50 µg/ml (A22). For treatments longer than 20 min cells were incubated in liquid culture with the respective antibiotic at the same concentration prior to imaging. The indicated incubation times denote the total incubation time in liquid and on the pad.

In experiments where we depleted MreCD and RodZ (strains EW07 and EW49 respectively) we grew cells in the presence of 50 or 100 µM IPTG respectively (EUROMEDEX #EU0008-B) and carbenicillin both during overnight growth and during regrowth as described above. We then washed the cells 3 times, diluted them and grew them in minimal medium in the absence of IPTG for 6 hours.

In experiments where we repressed the *mrdAB* operon (strain AV48/pcrRNA G20-R20/pKC128) we grew cells in the presence of kanamycin and chloramphenicol both during overnight growth and during regrowth as described above. We added anhydrotetracycline (aTc) (100 ng/ml, Acros Organic) upon dilution in minimal medium to induce the repression. We grew cells for 9 h prior to imaging both in the presence and absence of aTc.

In experiments where we imaged msfGFP-PBP2, TU230(attLHC943) cells are plated on and grown overnight in minimal media containing 25 µM IPTG. For the regrowth from saturated culture, we used minimal media containing the indicated IPTG concentration. We grew cells for 6, 8 or 10 hours to early exponential phase at 30°C. For the measurement of auto fluorescence, we grew MG1655 in the same way but without the addition of an antibiotic or IPTG.

In experiments where we imaged GFP-RodZ, we used 100 µM IPTG.

For MreB-curvature correlation measurements we grew NO53 or NO53/pDB192 overnight at 37°C in LB medium. We diluted the cultures by 1:500 and grew them at 30°C in LB medium for 2h. We then washed the cultures and diluted them 1:200 in MOPS rich medium (MOPS EZ Rich Defined Medium Kit. Cat.No. M2105, TEKnova) and grew them at 30°C for an additional 4h, such that the culture density remained below an OD600 of 0.1. The growth media of NO53/pDB192 cells contains carbenicillin at all times until the cells are harvested for microscopy. Cell division is inhibited by adding 1mM IPTG in liquid culture inducing *sulA* expression 30 min prior to imaging. Cells were placed on 1% agarose pads of the same media without any antibiotics but containing the same amount of IPTG for filamentation. Images were taken right after placing cells on a pad (NO53) or after maintaining cells for 30 min on the pad at the same 30°C (NO53/pDB192).

### 4. Microscopy

Single particle tracking of PAmCherry-PBP2 and msfGFP-PBP2 was performed on a custom-designed fluorescence microscope (here referred to as microscope 1) in TIRF (total internal reflection fluorescence) mode. The microscope was equipped with a 100x TIRF objective (Apo TIRF, 100x, NA 1.49, Nikon), three laser lines: 405 nm (Obis, Coherent), 488 nm (Sapphire, Coherent), 561 nm (Sapphire, Coherent), a dichroic beamsplitter (Di03-R488/561-t3-25x36, Semrock), an edge filter for PAmCherry imaging (BLP02-561R-25, Semrock), and a laser-line filter (NF561-18, Thorlabs). Shuttering of the 488 nm laser was controlled with an acusto optic tunable filter (AA Optoelectronics) or with shutters (Uniblitz, LS3 and TS6B, Vincent Associates). The 405 nm laser was controlled directly via an USB interface. Images were acquired with an EMCCD camera (iXon Ultra, Andor). All components were controlled and synchronized using µManager (Edelstein et al., 2010).

For high-frequency imaging with PAmCherry-PBP2, images were acquired with exposure time and intervals of 60 ms for a duration of 1 min. A weak UV-laser pulse (405 nm, 100-200 ms) was provided before the acquisition and every 200 frames during the measurements to activate new fluorophores. Low-frequency imaging was conducted with an exposure time of 1000 ms and with imaging intervals of 3.6 s or 1 s for a duration of 3.5 min or 3 min, respectively. We activated fluorophores once before the acquisition for 100-200 ms.

Single molecule tracking with cells carrying the msfGFP-PBP2 fusion requires a photo-bleaching phase prior to image acquisition. For high-frequency imaging, the sample was exposed to 488 nm laser in epi mode in order not to bias our analysis towards diffusive molecules as exposure to light in TIR mode would predominantly lead to a loss of bound molecules. After photo-bleaching, we immediately switched to TIR mode for image acquisition. Images are taken with 60 ms exposure time and intervals for a duration of 1 minute as for the PAmCherry fusion. For low-frequency imaging, the sample is first photo-bleached in TIR mode with high laser power. This step is followed by image acquisition while using a reduced laser power with an exposure time of 1000 ms and imaging intervals of 3.6 s for a duration of 4.5 minutes. Bleaching times for both cases were adjusted according to the PBP2 levels and it varies between 5 to 30 or 2 to 10 seconds for high and low frequency imaging respectively.

For measurements of cell shape, average fluorescence intensity, MreB-curvature correlations, and MreB rotation we used two inverted epi-fluorescence microscopes, an Eclipse Ti (Nikon) microscope (microscope 2) or a DeltaVision$^{TM}$ Elite (GE Healthcare) microscope (microscope 3). Microscope 2 is equipped with a 100x phase contrast objective (CFI PlanApo LambdaDM100X 1.4NA, Nikon), a solid-state light source (Spectra X, Lumencor Inc., Beaverton, OR), a multiband dichroic (69002bs, Chroma Technology Corp., Bellows Falls, VT), and excitation (485/25) and emission (535/50) filters. Images were acquired using a sCMOS camera (Orca Flash 4.0, Hamamatsu, Japan) with an effective pixel size of 65 nm. Microscope 3 is equipped with a 100x phase contrast objective (UPlanSApo 100X NA 1.4, Olympus), a multi-band dichroic beamsplitter (DAPI-FITC-mCh-Cy5) with excitation (475/28) and emission (525/48) filters, and a sCMOS camera (DV Elite, PCO-Edge 5.5) with an effective pixel size of 65nm.

For MG1655 auto fluorescence and msfGFP-PBP2 intensity measurements and for measurements of MreB-curvature correlations we used microscopes 2 after focusing on cells based on the phase-contrast signal.

For measuring MreB-msfGFP motion we first obtained phase-contrast images and then moved the focal plane 250 nm below the central plane of cells, in order to track MreB-msfGFP spots crossing the cell centerline. Images were taken every 1 s for a duration of 120 seconds.

### 5. Cell segmentation

Cell boundaries were detected from phase contrast microscopy images on microscope 2 or 3 using the MATLAB based cell segmentation tool Morphometrics (SimTK) (Ursell et al., 2017). The cell poles and the cell centerline were identified using the MicrobeTracker package (Sliusarenko et al., 2011). The spacing of subsequent points along the cell centerline was chosen as 0.5 pixels (32 nm). Cell width was measured as the median of all local widths.

A smoothened centerline ($x$- and $y$-coordinates Gauss-filtered, $\sigma$ = 3.5 steps) was used for a cell-internal orthogonal coordinate system, specifically to determine the local orientation of the cell.

### 6. PBP2 tracking

All images taken on microscope 1 were analyzed using custom Matlab code. First, we segmented bright field images using a semi-automated approach based on standard image processing tools to separate regions containing cells from background regions. For spatially separated cells this also allowed us to obtain a coordinate system for each cell.

PAmCherry-PBP2 spots in fluorescence images were identified as the local maxima of the denoised (bandpass filtered) images with intensity 3.5 times higher than background (bpass and pkfnd functions from https://site.physics.georgetown.edu/matlab/code.html). Sub-pixel resolution was achieved by finding the center of a two-dimensional Gaussian fitted to the intensity profile of each spot. We discarded peaks with a poor (residuals of fit) or broad (standard deviation of fit) Gaussian profile. Spots in subsequent frames were then connected into raw trajectories if their distance was below a threshold distance that is consistent with diffusion (high-frequency imaging) or persistent motion (low-frequency imaging) (Crocker and Grier, 1996). Threshold distances were 600 nm for imaging intervals of $\tau$ = 60 ms, 112.5 nm for $\tau$ = 1 s, and 225 nm for $\tau$ = 3.6 s. If tracking of a particle lead to a situation where a particle can be connected to more than one possible peak in the next frame we discontinued the tracking for this trajectory. We checked that low-frequency trajectories lie in cells by comparing fluorescent images and brightfield images.

msfGFP-PBP2 images from fast frequency imaging are analyzed in the same way except that peaks which have intensity 2 times higher than background are selected.

msfGFP-PBP2 spots from low frequency imaging were identified and tracked in the following way: Peaks were preselected as local maxima in the bandpass-filtered image (using a Laplacian of Gaussian filter with $\sigma$ = 1.8 pixels and pkfind, as above). We only considered local maxima in regions of the 2% highest intensity values that are 4-connected and contain at least 3 pixels each. We only consider peaks with nearest-neighbor distance higher than 3.5 pixels.

For further peak selection and sub-pixel localization we considered denoised raw images (using a Gauss filter with $\sigma$ = 0.4 pixels). We selected peaks with a ratio of peak signal to local background noise higher than 3. Local background intensity and noise are respectively the average and standard deviation of the values of the pixels at a distance included between 3 and 4 pixels of the center of the peak. To avoid very low intensity peaks we also discarded peaks with absolute signal (<2000 counts for 1s exposure, <1200 counts for 60ms exposure), and we remove the 1% of the peaks with highest intensities, as those likely represent clusters of molecules. Sub-pixel resolution was achieved by fitting a two-dimensional Gaussian to every selected peak. Peaks with standard deviation > 3 or with a ratio of the average absolute value of the residuals divided by peak intensity > 0.2 are removed.

Tracking was performed using the same code from (Crocker and Grier, 1996) with a maximum displacement between subsequent frames of 4 pixels (both for slow and fast tracking). For lifetime measurements with a time interval of 12 s, we increased to maximal displacement to 6 pixels.

### 7. MreB tracking

Fluorescence images were analyzed using a custom Matlab code: Images were first filtered in both space and time using a three-dimensional Savitzky-Golay filter with a filter size of 3 pixels in *xy*-directions time 3 points along the temporal dimension. Images were subsequently de-noised once more using a 2D-Gauss filter ($\sigma$ = 0.5 pixels).

Images were subsequently rescaled by a factor of 5 using spline interpolation to achieve sub-pixel resolution. MreB spots were detected as local maxima inside the cell boundary obtained by segmentation.

The local maxima were connected to construct raw trajectories based on their distance at consecutive time points (van Teeffelen et al., 2011) with a maximal displacement during subsequent time frames of 2 pixels.

### 8. Curvature analysis and MreB-curvature correlations

After obtaining the cell contours and the two poles of each cell we first computed the contour curvature all along the cell boundary by fitting a polygon to every contour point and its two neighboring points (MATLAB function LineCurvature2D, http://www.mathworks.com/matlabcentral/). Negative curvature values correspond to indentations and positive curvature values to bulges and poles. To eliminate the influence of noise the contour curvature $c$ is then obtained by Gauss-filtering the raw curvature values ($\sigma$ = 2 steps, corresponding to 65 nm).

To obtain MreB-msfGFP intensities along the cell boundary we first smoothened raw MreB-msfGFP images using a 2D-Gauss filter ($\sigma$ = 0.5 pixels). For interpolation of the GFP images at points close to the cell boundary we then corrected the contour coordinates $\boldsymbol{r}_i$ for systematic shifts between the phase-contrast-based cell contours and the GFP-intensity peaks corresponding to MreB filaments. Specifically, we obtained the corrected contour coordinates $\boldsymbol{r}_i^c$ from the Morphometrics-based contour coordinates $\boldsymbol{r}_i$ according to

$$\boldsymbol{r}_i^c = \boldsymbol{r}_i + \boldsymbol{\Delta r} - \boldsymbol{r}_{\perp i} s.$$

Here, the first term $\boldsymbol{\Delta r} = (-0.35, -0.78)$ pixels is a microscope-dependent displacement vector that accounts for a systematic shift between phase-contrast and GFP images. The second term shifts the contour by an amount $s$ = 140 nm inward (along a vector normal to the cell boundary $\boldsymbol{r}_{\perp i}$), such that the corrected contour passes through the positions of the GFP intensity peaks that correspond to the positions of MreB filaments on the sides of the cell. By convention, $\boldsymbol{r}_{\perp i}$ always points outward from the cell center. To make sure to take MreB-msfGFP peaks into account even if they are slightly displaced from the corrected boundary, GFP intensity values $I_i$ are then obtained by linear interpolation of the smoothened MreB-msfGFP image at the positions of the corrected contour and at two other points perpendicular to the boundary and spaced 0.5 pixel inward an outward:

$$I_i = \frac{1}{3}[I(\boldsymbol{r}_i^c - \boldsymbol{r}_{\perp i}\delta) + I(\boldsymbol{r}_i^c) + I(\boldsymbol{r}_i^c + \boldsymbol{r}_{\perp i}\delta)],$$

where $\delta$ = 65 nm. Subsequently, intensity values are normalized by the average taken over the full contour, the side walls, or the straight cell segments, respectively, depending on the analysis. To obtain the MreB enrichment as a function of curvature, the curvature values are binned and MreB intensities corresponding to those curvature vales are averaged. Subsequently, intensity values are normalized by the average intensity found close to zero curvature:

$$I_i^c = I_i / \langle I_i | -0.05/\mathrm{um} < c_i < 0.05/\mathrm{um} \rangle \ .$$

If a bin contains less than at least 0.1% of the data points per replicate, it is not displayed in the figure.

To remove the cell poles and potential septa from the analysis, we removed 2 um from each cell pole (according to the distance along the centerline) and 650 nm from the middle of the cell. The large distance from the poles was chosen, since the automated software does sometimes not identify the poles correctly. In those cases, contour points can fall into MreB-msfGFP-free regions at the poles even if the corresponding centerline points are more than just 0.5 um away from the automatically identified poles.

To determine correlations between MreB and contour curvature independently of spontaneous cell bending, we either concentrated on regions where the smoothened centerline was straight or we performed two independent normalization approaches:

To constrain our analysis to straight segments of the cell we considered only those boundary points corresponding to centerline curvatures with $|\kappa_i| < 0.05/\mu m$.

In the first normalization approach we renormalized the contour curvature by subtracting the contribution of cell bending according to

$$c_i^{\mathrm{corr}} = c_i \pm \kappa_i / \left(1 - \frac{\kappa_i w_i}{2}\right),$$

where $\kappa_i$ is the curvature of the smoothened centerline (*x*- and *y*-coordinates Gauss-filtered, $\sigma$ = 3.5 steps), and where $w_i$ is the local width of the cell. The centerline was smoothened to consider only the contribution of long-range bending rather than the impact of short-scale oscillations of boundary curvature. Positive/negative centerline-curvature values correspond to cells bent to the right/left along the direction of the centerline. The plus or minus signs correspond to the right or left side of the cell, respectively. The correction only works for segments of the cell with $\kappa_i < 0.5\, w_i$ to avoid divergence. For the cell segments excluding poles and potential septa this criterion was always fulfilled.

In the second normalization approach we renormalized the MreB intensity by the component expected due to cell bending:

$$I_i^{\mathrm{corr}} = I_i/(1 \pm \alpha\kappa_i w_i),$$

where alpha ~ 0.2 is a coefficient that accounts for the correlations observed between MreB intensity and smoothened centerline curvature (Fig. 7–SI Fig. 2D). As above, the plus/minus signs correspond to the inner/outer face of the bent cell, respectively.

### 9. Analysis of spots on cell boundaries and colocalization

To analyse the distribution of fluorescence peaks of cells expressing mCherry-PBP2, MreB-msfGFP, or GFP-RodZ and to perform colocalization measurements we first obtained fluorescence profiles on cell contours from epi-fluorescence images as described in Section 8.

Peak analysis: Fluorescence profiles were smoothened with a Gauss filter ($\sigma$ = 0.5 pixels). Subsequently, we subtracted the median intensity of every cell contour. Peaks were then detected as positive-valued local maxima (Figs. 3C, 6E).

The intensity-dependent peak density is the number of peaks with intensity *p* divided by total contour length of all cells $\sum_i^{N_{\mathrm{cells}}} l_i$ . For the total density of all peaks, we considered all peaks with peak height 3 times higher than the estimated intensity noise (gray regions in Figs. 3C, 6E). The noise was calculated in non-treated conditions. To that end, we filtered the raw fluorescence image with a 2D Gauss filter of $\sigma$ = 0.5 pixels and $\sigma$ = 3 pixels separately. The signal on the cell boundaries are then extracted from two sets of images. Next, we took the difference of signals belonging to same cells and calculated the average standard deviation of the differences, which is used as a readout for pixel noise.

The attenuation factor *S* is the fold change of the number of fusion proteins found in peaks between A22-treated and non-treated conditions. We calculated *S* as

$$S = T_{\mathrm{A22}}/T_{\mathrm{WT}}.$$

where

$$T_{\mathrm{WT/A22}} = \left(\sum_i^{N_{\mathrm{cells}}} \sum_j^{N_i} p_{ij}\right)/\sum_i^{N_{\mathrm{cells}}} l_i\ .$$

Here, $N_{\mathrm{cells}}$ is the number of cells, $N_i$ is the number of peaks in cell *I*, $p_{ij}$ is the intensity of peak *j* in cell *I*, and $l_i$ is the contour length of cell *i*.

For colocalization analysis, we extracted profiles as described above for two different fusions and calculate the Pearson correlation coefficient in each cell independently.

### 10. Determination of bound fraction and diffusion constant

We inferred bound fraction and diffusion constant from the distribution of effective diffusion constants, $D_{\mathrm{eff}}$, from single-molecule tracks. We calculated $D_{\mathrm{eff}}$ from the first 4 steps of each track, applying a linear fit of single-track MSD's $\langle x^2(t)\rangle$ according to $\langle x^2(t)\rangle = 4D_{\mathrm{eff}}t + 4\sigma^2$. The empirical distribution of $D_{\mathrm{eff}}$ was then compared with distributions generated from computational simulations. For both reference conditions of PAmCherry-PBP2 and msfGFP-PBP2 fusions (TKL130 and TU230(attLHC943) with 25 uM IPTG, respectively), we calculated distributions $p(D_{\mathrm{eff}}|D, \sigma)$ for diffusive molecules with different diffusion constant *D* and localization uncertainties $\sigma$. The combined distribution of a two-state population is then given by $p(D_{\mathrm{eff}}) = b\ p(D_{\mathrm{eff}}|D_{\mathrm{bound}} = 0, \sigma) + (1\text{-}b)\ p(D_{\mathrm{eff}}|D, \sigma)$, where we assume bound molecules to be immobile. For every combination [$D$, $\sigma$] we obtained the bound fraction *b* by fitting the peak value at $D_{\mathrm{eff}}$ = 0 µm$^2$/s to the experimental data. [$D$, $\sigma$] were then obtained by minimizing the residual sum of squares (RSS) between experimental and simulated distributions, considering values for $D_{eff}$ between -0.015 and 0.055 μm$^2$/s (see Fig. 1–SI Fig. 2A for PAmCherry-PBP2). We used a bin size of 0.005 μm$^2$/s.

For PAmCherry-PBP2 we varied $D$ between 0.015 and 0.06 μm$^2$/s with an interval of 0.005 μm$^2$/s and $\sigma$ between 0 and 40 nm with and interval of 5 nm. The parameter set of $D$ = 0.04 μm$^2$/s and $\sigma$ = 20 nm gave the lowest RSS. We confirmed that $D_{bound}$ = 0 μm$^2$/s by comparisons between distributions for zero and finite values of $D_{bound}$.

For the msfGFP-PBP2 fusion we obtained similar results: We varied $D$ between 0.0075 and 0.04 μm$^2$/s with an interval of 0.0025 μm$^2$/s and $\sigma$ between 0 and 50 nm with and interval of 5 nm, which led to $D$ = 0.01 μm$^2$/s, and $\sigma$ = 20 nm.

For all other datasets, we kept $\sigma$ = 20 nm fixed and only varied $D$ in simulations (between 0.0075 and 0.06 μm$^2$/s with an interval of 0.0025 μm$^2$/s) for each replicate to find the variation of the diffusion constant.

### 11. Velocity and orientation distributions of persistently moving PBP2 and MreB

Directed motion of individual trajectories can be inferred from a quadratic dependency of the single-particle mean squared displacement (MSD) on time according to

$$\langle x_i^2 \rangle = v_i^2 t^2 + b_i. \qquad (11.1)$$

Here, $v_i$ denotes the velocity of particle $i$ and $b_i$ is an offset reflecting the localization error.

To determine the velocity distribution of persistently moving PBP2 molecules or MreB filaments, respectively, we analyzed a subset of trajectories obtained with 1 s time intervals, for which each single-track MSD could be well fit by a quadratic function of the form of Eq. (11.1) according to the coefficient of determination $R^2$. We considered trajectories with at least 4 time steps. For PBP2 we considered trajectories with a maximum of ten time steps to prevent a bias towards slowly moving molecules that are more likely to remain in the TIR field of illumination for long times. MreB trajectories are longer in time than PBP2 trajectories, since they are obtained from filaments containing many msfGFP molecules, and they are obtained from a larger field of view since their movement is measured in epi-fluorescence mode. For comparability with PBP2 tracks we therefore constrained the MreB trajectories by analyzing the 10 displacement steps around the point closest to the cell centerline. These displacements are found within a field of view also detectable in TIRF mode.

The velocity distributions for different minimal $R^2$ values are plotted in Fig. 1–SI Fig. 5D. As expected, low velocities corresponding to immobile molecules contribute less with increasing minimal $R^2$. Accordingly, both MreB and PBP2 showed a mildly increasing average velocity as a function of the minimum $R^2$ (Fig. 1 SI Fig. 5D). For comparison between PBP2 and MreB in Fig. 1F-G we generated velocity distributions from tracks that showed $R^2$ values >= 0.9, corresponding to ~10% of PBP2 tracks that showed the highest $R^2$ values (Fig. 1 SI Fig. 5C), yielding average velocities of 18.3 nm/s for PAmCherry-PBP2 and 16.5 nm/s for MreB-msfGFP filaments (Fig. 1 SI Fig. 5D).

Performing the same analysis on PAmCherry-PBP2 trajectories generated with a time interval of 3.6 s we obtained an average velocity of $v$ = 14.2 nm/s for persistent tracks ($R^2$ >= 0.9). The average velocity is lower than for the 1 s dataset because fast-moving molecules are observed less often than slowly moving molecules, as they leave the TIR field of view at higher probability (Fig. 1 SI Fig. 5E-F). For msfGFP-PBP2 we obtained an average velocity of 13.5 nm/s.

To obtain the orientation of persistent PBP2 tracks we calculated the angle between the end-to-end vector of persistent trajectories measured with time interval of 3.6 s (same criteria of $R^2$ >= 0.9 and a minimum length of 4 time points) and the cell orientation. For MreB we calculated the orientation in the same way and on the same data for which we obtained the velocity distribution.

### 12. Localization accuracy in low-frequency imaging

We determined the experimental localization accuracy from the distribution of displacements from measurements at 3.6 s time intervals for PAmCherry-PBP2 (Fig. 1 SI Fig. 6A). We restricted our analysis to immobile molecules according to a simple criterion (end-to-end distance less than 200 nm for tracks of at least 7 steps). The standard deviation $\sigma_d$ of a Gaussian fit to the distribution of displacements in a single spatial direction is determined by $\sigma_d^2 = \langle (x_i - x_{i+1})^2 \rangle = 2\sigma^2$, where $\sigma$ is the localization uncertainty. Therefore, $\sigma = \sigma_d/\sqrt{2}$ . For PAmCherry-PBP2 $\sigma = 25\mathrm{nm}$.

### 13. Simulation of persistent and immobile tracks

To establish a criterion for reliably classifying persistently moving and immobile states in experimental PBP2 trajectories, we computationally simulated tracks of immobile or persistently moving molecules resembling the tracks observed by microscopy using 3.6 s time intervals. To that end we randomly picked a trajectory length (number of steps *n*) from an exponential distribution with $\langle n \rangle = 3.5$ that resembled the experimental length distribution (Fig. 1 SI Fig. 6B). For the simulation of persistently moving or immobile molecules we imposed a constant step size in the *x*-direction corresponding to the experimental velocity (*v* = 14 nm/s for PAmCherry-PBP2) for persistent molecules or to *v* = 0 nm for immobile molecules. To account for the localization uncertainty we subsequently added to all *x*- and *y*-coordinates a random displacement drawn from a normal distribution with a mean of 0 nm and a standard deviation equal to the localization accuracy for low-frequency imaging 25 nm (for PAmCherry-PBP2).

Trajectories with transitions between immobile and persistent states were obtained by randomly selecting sub-trajectories from a single 1000-step long trajectory containing transitions between persistent and immobile states. Transition rates were obtained from experimental data (see next paragraph).

### 14. Determination of persistent and immobile states and switching rates

For each time point we calculated the smoothed local velocity by dividing the displacement during the surrounding *w* time steps by the time lag *wτ*:

$$v(t) = \boldsymbol{r}\left(t + \frac{w}{2}\right) - \boldsymbol{r}\left(t - \frac{w}{2}\right)/w\tau,$$

Here $\boldsymbol{r}(t) = \big(x(t), y(t)\big)$ is the position at time *t*, and $\tau$ = 3.6 s is the imaging time interval. We classified the particle as either immobile or persistently moving at time *t* if *v* was smaller or bigger than the threshold velocity $v_{thr}$, respectively.

We applied different values for *w* and $v_{thr}$ to simulated trajectories of PAmCherry-PBP2 (*v* = 14 nm/s; sigma = 25 nm) to find the parameter combination that reliably detected dynamic states in simulated tracks with more than 99% success rate (Fig. 1 SI Fig. 3C). We chose a window size of *w* = 4 and a velocity threshold of $v_{thr}$ = 8 nm/s (classifying 99.6% of segments of simulated immobile molecules as immobile and 99.6% of simulated persistent molecules as persistent) (Fig. 1 SI Fig. 6C). Since we found almost identical average velocity for msfGFP-PBP2, we used the same window size and velocity threshold for persistence classification.

We calculated transition rates $k_{ip}$ or $k_{pi}$ by counting the number of transitions from immobile to persistent or persistent to immobile states, respectively, and dividing by the total duration of persistent or immobile states observed. Here, we ignored intermittent segments of duration of a single time step.

All error bars denote standard errors between replicates.

### 15. Testing the two-state model of immobile and persistent states

To test whether the dynamics of PBP2 molecules is compatible with a model of molecules residing in either of two possible states we measured single-particle MSD's of track segments identified as either immobile or persistent (Fig. 1–SI Fig. 6A). The average MSD of segments classified as persistently moving increased quadratically with time, while the average MSD of segments classified as immobile remained nearly constant (Fig. 1 SI Fig. 6 D-E).

To test whether deviations of single-particle MSDs from the average were due to transitions between the two states we analyzed computationally simulated tracks of our two-state model containing transitions. We then used the same classification criterion and MSD analysis on simulated tracks as on experimental tracks. For simulations of persistent segments we adjusted the velocity of $v$ = 12 nm/s (compared to 14 nm/s above) that yielded better agreement of the average MSD curves of persistent segments with experiments (Fig. 1–SI Fig. 6D). The difference is likely due to the fact that we consider here all persistent segments while above we obtained the velocity from the 10% most persistent full tracks according to the $R^2$-criterion.

Simulations and experiments showed very similar distributions of single-particle MSD's (Fig. 1–SI Fig. 6F), suggesting that bound PBP2 molecules are indeed either immobile or persistently moving, but not found in a qualitatively different slowly-moving state.

### 16. Calculation of the unbinding rate based on fluorescence-lifetime measurements

To obtain the unbinding rate $k_{bd}$ in non-treated or A22-treated cells we measured lifetime distributions of tracks $f(n,\tau)$ obtained with 1 s exposure time and different time intervals $\tau$ = 1 s or 12 s (Fig. 2). Here, $n$ is the number of steps a track is observed corresponding to the lifetime $t = n\tau$. At first, we assume two random processes to contribute to particle loss: GFP bleaching with a probability $p_b$ per time frame, and a second process with an apparent track termination rate $k_a$, corresponding to a termination probability $p_a = 1 - \exp[-k_a\tau]$.

For A22-treated cells, $k_a$ is caused by unbinding only, that is, $k_{bd} = k_a$. For non-A22-treated cells, $k_a$ subsumes unbinding and particles leaving the field of view due to persistent motion. While the probability of molecules leaving the field of view is not independent of track duration, this assumption does not affect our calculation of $k_{bd}$, as we will see below.

For both conditions (-A22, +A22), we simultaneously fit the two lifetime distributions to the above model, considering tracks between 3 to 7 steps (4-8 localizations). For A22-treated cells we obtained $p_b = 0.43 \pm 0.08$ and $k_a = k_{bd} = 0.021 \pm 0.008\ s^{-1}$. The unbinding rate corresponds to an average lifetime of the bound state of 48 ± 18 s.

For non-A22-treated cells we obtained $p_b = 0.39 \pm 0.08$ and $k_a = 0.035 \pm 0.007\ s^{-1}$. To estimate the contribution of persistent motion to the apparent unbinding rate, we then conducted simulations of bound molecules transitioning between persistent and immobile states: Molecules started from random positions within the field of view (width 600 nm) either moving persistently with speed of $v = \pm$ 14 nm/s perpendicular to the central axis, or resting immobile. Transitions between the two states occurred at experimental rates $k_{ip}$ and $k_{pi}$, respectively. The probability of bleaching was set equal to $p_b$. In a second set of simulations, we changed either of the two rates to maintain the measured persistent fraction of $p$ = 80%, that is, either $k_{ip}^{corr} = k_{pi}\, p/(1-p)$ or $k_{pi}^{corr} = k_{ip}\,(1-p)/p$ (Fig. 2–SI Fig. 1). We then measured track-length distributions from simulations with different unbinding rates $k_{bd}$ to infer the range of unbinding rates $k_{bd}$ compatible with the experimentally obtained apparent termination rate $k_a$ (Fig. 2–SI Fig. 1).

In both conditions, experimental lifetime distributions of 1 s data are dominated by bleaching. Therefore, the time-dependent process of persistent molecules leaving the field of view only affects the 12 s distributions. While those distributions are not perfectly exponential, we could still fit them by an exponential function over the window of 3 to 7 steps, considered here.

### 17. Measuring transitions from diffusive to bound states (Bound-Molecule FRAP)

In order to determine the transition rate from the diffusive to the bound state $k_{db}$ we measured the bound fraction at different time points after bleaching the field of view, conceptually similarly to classical FRAP (fluorescence recovery after photobleaching) experiments. In a first step we aimed to activate all PAmCherry fluorophores in the TIRF field of view with a 1.5 s exposure of 10-fold increased UV intensity compared to our standard protocol (see above). We introduced a waiting time of 2 seconds in the dark in order to let diffusive photo activated PBP2 molecules escape the field of view. Then, we photo bleached for 4 seconds with normal excitation intensity. After a recovery period of 0-10 minutes in the dark, we acquired images in high-frequency mode without any additional photo activation for a duration of 48 seconds. In this way, we were able detect PAmCherry-PBP2 molecules, which were activated in the first step, which were able to escape the field of view during the 2 second pause, and which then reentered the field of view, where they either remained diffusive or bound to their substrate.

### 18. Quantification of expression level from fluorescence

For PAmCherry-PBP2 we counted the number of fluorescent spots observed per cell after a single activation pulse of the UV laser. For the comparison between native and overexpression levels see Fig. 1 – SI Fig. 1D.

We quantified the different levels of msfGFP-PBP2 obtained on microscope 2 (see above) by measuring the total GFP fluorescence intensity per cell, where cell outlines were obtained by segmentation of phase-contrast images using Morphometrics (SimTK) (Ursell et al., 2017). We subtracted contributions from auto-fluorescence per pixel as obtained from imaging wildtype *E. coli* (MG1655) cells. For a comparison between different induction levels see Fig. 1 – SI Fig. 7B.

### 19. Model of MreB surface area fraction

To estimate the fraction of the cytoplasmic membrane covered by MreB filaments we assume for simplicity that all MreB proteins of the cell are part of dimers of MreB protofilaments (Salje et al., 2011). A substantial fraction of proteins is found as cytoplasmic monomers. Our estimate is therefore rather conservative. Previous measurements suggest that there is a broad distribution of filament lengths with many filaments as long as 1 um (Ouzounov et al., 2016). As a conservative estimate we assume that all filaments are 100 nm in length and have a repeat length of 5 nm (Salje et al., 2011). Each of the idealized filaments therefore contains 40 monomers. With an average of 2000 or 11000 proteins per cell in poor or rich growth media, respectively (Li et al., 2014) the cytoplasmic membrane is decorated with up to 200 filaments according to this simple model. Cells grown in our conditions have a surface area of about 9 $um^2$. Assuming that interactions between PBP2 and MreB require PBP2 to be within a distance to MreB of 5 nm, MreB filaments occupy about 2 % of the area of the cytoplasmic membrane.

### 20. Model to test the contribution of diffusing PBP2 molecules to rod-complex activity

Cross-links with neighboring glycan strands are formed every other di-sugar subunit. Each subunit is about 1 nm long (Boal and Boal, 2012). Thus, the rate of transpeptidation corresponding to a high but common speed of MreB of 30 nm/s is $\lambda$ = 15/s. Lee *et al.* argued that after forming one cross link, PBP2 would detach and diffuse in the cell envelope to find a new site for cell-wall cross-linking (Lee et al., 2014).

The number of PBP2 enzymes in the cell is about between 100-300 in nutrient-rich medium and 60-75 in poor medium according to radiolabeling (Dougherty et al., 1996) or ribosome profiling (Li et al., 2014). We thus wondered whether free diffusion of such a small number of enzymes could account for the experimentally observed rate of cross-link formation, or whether free diffusion would limit this process. Alternatively, we also considered that molecules underwent facilitated diffusion along one-dimensional tracks such as the cytoskeleton MreB (Oswald et al., 2016), similarly to the phenomenon of transcription factors searching their target on chromosomal DNA (Mirny et al., 2009).

We conducted overdamped Brownian-dynamics simulations of $N$ = 100 enzymes [interpolating between measurements made for poor and rich media (Dougherty et al., 1996; Li et al., 2014)] in a rectangular domain of 3 x 3 um with periodic boundary conditions in $x$- and $y$-directions, thus approximating the cylindrical surface of a rod-like *E. coli* bacterium of 1 um width and 3 um length (Fig. 1 – SI Fig. 1B) and ignoring the shape of the cell poles.

The overdamped Brownian motion of PBP2 in our model is governed by the Langevin equation for its position

$$\dot{\boldsymbol{r}} = D\boldsymbol{\zeta}$$

where the dot denotes a time derivative, $D$ = 0.06 um$^2$/s is the experimental diffusion constant, and $\boldsymbol{\zeta}$ is the zero mean Gaussian white noise random displacement originating from the solvent. Its variance is given by

$$\overline{\boldsymbol{\zeta}(t)\boldsymbol{\zeta}(t')} = 2\delta_{ij}\delta(t - t'), i, j = x, y,$$

where the bar denote a noise average.

A number of $n$ = 10 circular cross-linking sites of diameter $a$ = 10 nm are placed at random locations in the rectangular domain. Once a diffusing molecule hits any of the cross-linking sites, a cross-linking event is registered to occur. Note, that this model is based on the conservative estimate that every encounter between enzyme and cross-linking site leads to a successful reaction. To prevent rapid return of an enzyme to the same site we introduce a deterministic latency time $t_{off}$ after every encounter during which an enzyme can diffuse but not facilitate a reaction. This latency time could reflect the typical time it takes to conduct one reaction or a combination of different microscopic effects. The reaction rate per site $\gamma$ is then calculated as the mean number of enzyme-site encounters per site per total simulated time.

We consider latency times larger than 0.1 ms (Fig. 8B), a time that is needed for a PBP2 enzyme to explore an area similar to the size of an enzyme (5 nm). In this regime, the encounter rate depends only weakly on $t_{off}$ (Fig. 8B). Thus, only a minor fraction of enzymes re-encounters the same site shortly after leaving it (Fig. 8C). Notably, the effect of $t_{off}$ on rebinding is much weaker than in the previously studied cases of finding a membrane receptor from the cytoplasm or of binding a receptor in the 3D bulk (Mugler et al., 2012), where the probability of rebinding decays algebraically with the latency time for short times and exponentially for long times. Results are nearly independent of target numbers $n$ (not shown).

We next considered the possibility that PBP2 undergoes facilitated diffusion along one-dimensional tracks, such as MreB filaments: MreB forms circumferentially oriented filaments of up to 1 um in length (Ouzounov et al., 2016). PBP2 enzymes and other cell-wall proteins interact with MreB filaments (Kruse et al., 2004; Morgenstein et al., 2015), and PBP2 was observed to partially co-localize with MreB filaments (Lee et al., 2014). To test the possible influence of linear tracks on the encounter rate we extended the model introduced above by adding unidirectional filaments of length $l$ to every rod-complex site (filaments are oriented along the $y$-axis). PBP2 molecules cannot cross filaments. Instead, a PBP2 molecules that encounters a filament, diffuses along the filament with the same diffusion constant $D$ until it either a) hits the reaction site, b) reaches one of the two filament ends and returns to 2D diffusion, or c) is randomly displaced from the filament by an amount $\Delta x = 2a$ with rate $k_{\mathrm{off}}$. Only after hitting the target is an enzyme inactive for the latency time $t_{\mathrm{off}}$.

### 21. PBP2 Mass-spectrometry

To quantify relative changes of protein levels between conditions and strains, we used Data Independent Acquisitions (DIA) following (Bruderer et al., 2017). For absolute quantification of PBP2 levels, we used a targeted proteomics approach by Parallel Reaction Monitoring (PRM) (Bourmaud et al., 2016; Gallien et al., 2012; Peterson et al., 2012).

#### Preparation of *E. coli* whole protein extracts

Cells were collected by centrifugation (4000g, 10 minutes at 4°C) around $OD_{600}$ 0.15. For absolute quantification of PBP2, an aliquot part was taken from each culture in order to determine cell number by colony counting. Supernatant was removed and cell pellets were flash-freeze in liquid nitrogen and stored at -80°C. Cells were suspended in 250 µl of Urea buffer 8M (Sigma U4883). Cooled cells were lysed by sonication (Fisherbrand FB120) (alternating 3 cycles of 30 seconds ON with 40% amplitude and 15 seconds OFF to cool down the sample). Protein concentration was determined using a Bradford-based colorimetric assay (Bio-Rad 5000006) (Bradford, 1976) with known concentrations of bovine serum albumin (Sigma) as a standard. Proteins samples were diluted with 2x phosphate buffered saline (PBS) in order to decrease Urea concentration and be compatible with the colorimetric assay. For the quantification of absolute numbers of PBP2 we used colony counting and measured protein concentration led, which resulted in an average of 105 fg of proteins per cell.

Digestion of proteins

All protein samples were denatured in 8 M urea in Tris HCl 100 mM pH 8.0. Proteins disulfide bonds were reduced with 5 mM tris (2-carboxyethyl)phosphine (TCEP) for 20 min at 23° C and further alkylated with 20 mM iodoacetamide for 30 min at room temperature in the dark. Subsequently, LysC (Promega) was added for the first digestion step (protein to Lys-C ratio = 80:1) for 3 h at 30° C. Then the sample was diluted to 1 M urea with 100 mM Tris pH 8.0, and trypsin (Promega) was added to the sample at a ratio of 50:1(w/w) of protein to enzyme for 8 h at 37° C. Proteolysis was stopped by adding 1% formic acid (FA). Resulting peptides were desalted using Sep-Pak SPE cartridge (Waters) according to manufacturer instructions. Peptides elution was done using a 50% acetonitrile (ACN), 0.1% FA buffer. Eluted peptides were lyophilized and then stored until use.

For Data Independent Acquisitions (DIA) and Parallel Reaction Monitoring (PRM) (see below), iRT peptides (Biognosys) were spiked into all samples as recommended by manufacturer.

Peptide Fractionation for spectral library

Peptide fractionation was done using poly(styrenedivinylbenzene) reverse phase sulfonate (SDB-RPS) stage-tips method as described in (Kulak et al., 2014; Rappsilber et al., 2007). Briefly, 3 SDB-RPS Empore discs were stacked on a P200 tip and used to fractionate 30 µg of peptides. Four serial elutions were applied as following: elution 1 (80mM Ammonium formate, 20% (v/v) ACN, 0.5% (v/v) FA), elution 2 (110mM Ammonium formate, 35% (v/v) ACN, 0.5% (v/v) FA), elution 3 (150mM Ammonium formate, 50% (v/v) ACN, 0.5% (v/v) FA) and elution 4 (80% (v/v) ACN, 5 % (v/v) ammonium hydroxide).

All fractions were dried and resuspended in 0.1% formic acid before injection. For all fractions, iRT peptides were spiked as recommended by Biognosys.

#### LC-MS data acquisitions

Data Independent Acquisitions (DIA)

LC-MS/SM analysis of digested peptides was performed on an Orbitrap Q Exactive HF mass spectrometer (Thermo Fisher Scientific, Bremen) coupled to an EASY-nLC 1200 (Thermo Fisher Scientific). Peptides were loaded and separated at 250 nl/min on a home-made C18 50 cm capillary column picotip silica emitter tip (75 µm diameter filled with 1.9 µm Reprosil-Pur Basic C18-HD resin, (Dr. Maisch GmbH, Ammerbuch-Entringen, Germany)) equilibrated in solvent A (2% ACN, 0.1 % FA). Peptides were eluted using a gradient of solvent B (80% ACN ,0.1 % FA) from 3% to 6% in 5 min, 6% to 29% in 130 min, 29% to 56% in 26 min, 56% to 90% in 5 min (total length of the chromatographic run was 180 min including high ACN level steps and column regeneration). Mass spectra were acquired in data-independent acquisition mode with the XCalibur 4.1.31.9 software (Thermo Fisher Scientific, Bremen).

Each cycle was built up as follows: one full MS scan at resolution 30 000 (scan range between 400 and 1200 m/z), AGC was set at $3*10^6$, ion trap was set at 50 ms. All MS1 was followed by 40 isolation windows of 20 m/z, covering the MS1 range from 400 m/z to 1200 m/z. The AGC target was $2*10^5$, and NCE was set to 27. All acquisitions were done in positive and profile mode.

Parallel Reaction Monitoring acquisitions for absolute quantification of PBP2 (PRM)

Peptides chosen and used for absolute quantification of PBP2 were based on the FASTA sequence obtained from UniprotKB database (The Uniprot Consortium, 2015) and MS evidence of identification. Peptides sequences are SGTAQVFGLK and VDNVQQTLDALR (Aqua UltimateHeavy, Thermo Fisher Scientific). Targeted peptides and their heavy forms were imported into Skyline (MacLean et al., 2010) to generate precursor ion inclusion list that also contained instrument control parameters for Xcalibur to detect peptides using PRM-MS. Information on iRT peptides (Biognosys) were also generated.

Heavy peptides synthetized from PBP2 sequence were spiked at $16 fmol.µl^{-1}$ in each sample. Each sample was injected at a known concentration with iRT peptides (as recommended by Biognosys) and 50 fmol of heavy peptides. Quantity of peptide injected on column was controlled by UV absorbance at 280 nm and tryptophan absorbance.

PRM was performed on an Orbitrap Q Exactive HF mass spectrometer (Thermo Fisher Scientific, Bremen) coupled to an EASY-nLC 1200 (Thermo Fisher Scientific). Peptides were loaded and separated at 250 $nl.min^{-1}$ on a home-made C18 50 cm capillary column picotip silica emitter tip (75 µm diameter filled with 1.9 µm Reprosil-Pur Basic C18-HD resin, (Dr. Maisch GmbH, Ammerbuch-Entringen, Germany) equilibrated in solvent A (2% ACN, 0.1 % FA). Peptides were eluted using a gradient of solvent B (80% ACN, 0.1 % FA) from 5% to 10% in 1 min, 10% to 30% in 82 min, 30% to 50% in 5 min, 50% to 95% in 5 min (total length of the chromatographic run was 105 min including high ACN level steps and column regeneration). Mass spectra were acquired XCalibur 4.1.31.9 software (Thermo Fisher Scientific, Bremen). The acquisition method combined a full scan method with a time-scheduled sequential PRM method. For the full MS, a scan range of 350 to 1500m/z, an orbitrap resolution of 60000, and an AGC value of a $3*10^6$ were used. An orbitrap resolution of 60000, a maximum IT set at 110 ms, an isolation window selection of 1.2 m/z, AGC target was $2*10^5$ and NCE fixed at 28 were used. Targeted, heavy and retention time peptides (iRT peptides, Biognosys) were listed in an inclusion list and monitored.

### Data analysis

Data Analysis for spectrum library building and DDA analysis of Co-IP

For spectral library purposes and DDA experiments, MaxQuant (Tyanova et al., 2016a) 1.5.5.3 was used. Raw data were analyzed against an *E. coli* database (6071 entries, downloaded from Uniprot on 10/03/2016).

The following search parameters were applied: carbamidomethylation of cysteines was set as a fixed modification, oxidation of methionine and protein N-terminal acetylation were set as variable modifications. The mass tolerances in MS and MS/MS were set to 5 ppm and 20 ppm respectively. Maximum peptide charge was set to 7 and 7 amino acids were required as minimum peptide length. A false discovery rate of 1% was set up for both protein and peptide levels.

Data analysis was done mainly using Excel and Perseus environment (Tyanova et al., 2016b).

Data analysis for DIA acquisitions

DIA experiments were analyzed using Spectronaut X (v. 11 Biognosys AG). Dynamic mass tolerance at the MS1 and MS2 levels was employed. The XIC RT Extraction Window was set to Dynamic with a correction factor of 1. Calibration mode was set to automatic with nonlinear iRT calibration and precision iRT enabled. Decoys were generated using the scrambled method and a dynamic limit (default settings). P value estimation was performed using a kernel density estimator. Interference correction was enabled with no proteotypicity filter. Major grouping was by Protein-Group ID, and minor grouping was by stripped sequence. The major group quantity was mean peptide quantity. The major group top N was enabled with a minimum of 1 and a maximum of 3. Minor group quantity was mean precursor quantity. The minor group top N was enabled with a minimum of 1 and a maximum of 3. The quantity MS-Level was MS2, and quantity type was area. Q value was used for data filtering. Cross run normalization was enabled with Q value sparse row selection and local normalization. The default labeling type was label-free with no profiling strategy and unify peptide peaks not enabled. The protein inference workflow was set to automatic.

PRM data analysis

Raw mass spectrometry data were exported to Skyline-daily (version 4.1.1.18179) for identification of transitions and peak area integration. Data were exported in .csv file format and analyzed in Excel.

### 22. Strain list

| Strain | Genotype | Source |
|---|---|---|
| TKL130 | MG1655 *mrdA*::*PAmCherry-mrdA* | (Lee et al., 2014) |
| EW07 | TKL130 Δ*mreCD*, pFB121 | This study |
| EW49 | TKL130 Δ*rodZ*, pFB290 | This study |
| AV48 | *186attB*::*Ptet-dcas9*, *mrdA*::*rPAmCherry-mrdA*, pKC128 | This study |
| NO53 | MG1655 *mreB-msfGFP*$^{sw}$ (after removal of KAN resistance cassette from NO50) | (Ouzounov et al., 2016) |
| TU230(attLHC943) | MG1655 *mrdA*::*aph* ($P_{lac}$::*msfgfp-mrdA*) | (Rohs et al., 2018) |
| TU230(attLPR122) | MG1655 *mrdA*::*aph* ($P_{lac}$::*msfgfp-pbpA*(L61R)) | (Rohs et al., 2018) |
| S352 | MG1655 Δ*ponA*::*aph* | This study |
| FB83 | MG1655, *lacIZYA*::frt, *mreB-mCherry*$^{SW}$ *yhdE*::frt | (Bendezú and Boer, 2008) |
| AV07 | MG1655 *mrdA*::*mcherry-mrdA* | (Vigouroux et al. 2018) |
| FB60 (iFB273) | MG1655 *lacIZYA*::*frt*, *rodZ*::*aph*, *Plac*::*gfp-rodZ* | (Bendezú and Boer, 2008) |
| S504 | FB83, *rodZ*::*aph*, $P_{lac}$::*gfp-rodZ* | This study |
| S505 | AV07, *rodZ*::*aph*, $P_{lac}$::*gfp-rodZ* | This study |

### 23. Plasmid list

| Plasmid | Relevant genotype | Source |
|---|---|---|
| pKC128 | *cat* $P_{mrdA}$::*rPAmCherry-mrdA* | (Lee et al., 2014) |
| pFB121 | *bla lacI*$^{q}$ $P_{lac}$::*mreC mreD* | (Bendezú and Boer, 2008) |
| pFB290 | *bla* $P_{lac}$::*rodZ* | (Bendezú et al., 2009) |
| pDB192 | *bla* $P_{lac}$::*sfiA* | (de Boer et al., 1990) |
| pcrRNA G20-R20 | CRISPR array cloning vector | (Vigouroux et al. 2018) |
| pTKred | *aadA* λ-Red enzymes | (Kuhlman and Cox, 2010) |
| pAV10 | PPhlF-*Cas9*-gRNA | (Vigouroux et al., 2018) |

| | | |
|---|---|---|
| pKD13 | Template vector for gene deletion. | (Datsenko and Wanner, 2000) |

### 24. Primer list

| **Primer** | **Sequence** |
|---|---|
| DmreC_fw | ATCGGATGCAGGCAGGGGAAGTGTCTGTTTACCCTGCCTGGTC TGATACGATAAGTGTAGGCTGGAGCTGCTTC |
| DmreC_rv | TCAGCAAGAAAATCCACGGCCAGAGCACCCCATTGACTACACT ACTCCAGAATTCCGGGGATCCGTCGACC |
